# Postnatal Pax7-expressing limb cells are multipotent and generate non-myogenic lineages that persist into adulthood

**DOI:** 10.64898/2026.04.21.719595

**Authors:** Stamatia Gioftsidi, Takuto Hayashi, Sidy Fall, Sabrina Jagot, Matthieu Dos Santos, Fabien Le Grand, Frederic Relaix, Philippos Mourikis

## Abstract

Organs are composed of a complex arrangement of diverse cell types that can originate from independent cell lineages or shared progenitors. Skeletal muscle is derived from mesodermal Pax7^+^ stem/progenitor cells that can differentiate into myoblasts to form muscle fibers. During embryogenesis, however, somitic Pax7^+^ cells can also give rise to non-muscle cell types, including dermis and adipocytes. Here, we asked whether Pax7^+^ cells retain such multipotency during early postnatal growth of limb muscles. Using lineage tracing, we uncovered unexpected plasticity at early postnatal days, leading to the generation of multiple non-myogenic lineages, including a previously unrecognized Pax7-derived subpopulation of fibro-adipogenic progenitors that we termed Pax7^FAPs^ and further investigated. Using mouse models, we further show that Notch signaling primes neonatal Pax7^+^ cells toward a fibrogenic molecular identity at the expense of myogenic differentiation, thereby biasing their trajectory toward a fibrogenic fate. In the adult muscle, long-term tracing revealed that neonatally produced Pax7^FAPs^ persist into adulthood. In addition, injury in adult muscle triggered de novo generation of Pax7^FAPs^, which displayed higher proliferative capacity than resident stromal cells. This newfound multipotency of postnatal Pax7^+^ cells adds a new dimension to our understanding of cellular contributions during postnatal muscle development and regeneration.

## Introduction

Skeletal muscle formation in vertebrates is marked by three distinct developmental waves that define the embryonic, fetal and postnatal myogenesis ^1^. In the trunk, skeletal muscles arise from the dorsal domain of the somites, the dermomyotome, which harbors the founder muscle stem cells (MuSCs). Embryonic MuSCs are marked first by PAX3 and later by PAX7, paralogues of paired homeobox transcription factors, and ultimately give rise to the adult MuSCs ^2-4^. Lineage experiments on embryonic and fetal *Pax7*^*+*^ cells, however, have shown tracing into non-myogenic lineages. Using a tamoxifen inducible *Pax7*^*CreERT2*^ allele for genetic labeling at embryonic day (E) 9.5 revealed the prominent contribution of *Pax7*^*+*^ cells to skeletal muscle, brown adipose tissue, and dorsal dermis. However, by E12.5, the contribution became restricted to muscle ^5^. Subsequent single cell RNA-seq analysis at E12.5, E14.5 and E16.5 on cells from the trunk of *Pax7*^*CreERT2*^*;Rosa26*^*YFP*^ embryos induced with tamoxifen at E9.5, confirmed the generation of diverse cell types by progenitors during mid-gestation ^6^. These included adipose and dermis cells, which likely derived from somitic mesoderm, whereas the identification of neuronal cells was likely the outcome of *Pax7* expression in the dorsal neural tube ^6^. Focusing on the extraocular muscles, Grimaldi and colleagues (7) identified two embryonic populations marked by *Myf5*^*Cre*^, one with a myogenic, *Myod*^*+*^*/Pdgfa*^+^ signature and one with a mesenchymal signature that was positive for the receptor *Pdgfra*. The *Pdgfra*^*+*^ descendants of *Myf5* cells were found in a region of mesoderm-derived cells and away from neural crest-derived connective tissue. Thus, the existence of an embryonic bipotent progenitor was proposed that gives rise to muscle and connective tissue cells in muscles lacking neural crest ^7^. Of interest, a reciprocal mechanism was uncovered at the level of the myotendinous junction (MTJ), where lateral plate derived fibroblasts were shown to contribute directly to the muscle lineage ^8,9^. Recently, a *pax7*+/*pdgfra*+ population of myoblasts was described in post-larval, juvenile rainbow trout, which was associated with the switch from hyperplastic to hypertrophic muscle growth ^10^.

Within adult skeletal muscle, PAX7 expression is restricted to MuSCs. These cells are indispensable for muscle regeneration and behave predominantly as unipotent stem cells, generating myofibers and self-renewing to maintain the MuSC pool ^11-14^. In dystrophic muscle, however, MuSCs aberrantly acquire a fibrogenic fate via a TGFb-mediated pathway, thus potentially contributing to the establishment of fibrosis ^15,16^. Despite this contribution, the main source of pathological fibrosis and fatty accumulation in dystrophic muscle is the mesenchymal progenitors that lay in the interstitial space between myofibers. These cells are collectively known as fibro-adipogenic progenitors (FAPs), they express PDGFRa and, in healthy muscle, are indispensable for steady-state muscle maintenance and regeneration ^17,18^.

Embryonic stem cells in diverse tissues are multipotent and progressively differentiate into lineage-restricted unipotent precursors that fuel postnatal development ^19^. However, in the skeletal muscle, it remains unclear when and whether Pax7^+^ cells adopt a unique myogenic fate. Here, we combined *in vivo* lineage tracing and single nucleus RNA-seq (snRNA-seq) to map the cellular trajectories of the Pax7^+^ stem/progenitor cells in the limbs during the first three weeks of postnatal development and to track their long-term fates into adulthood. Unexpectedly, we identified rare non-myogenic populations arising from the Pax7 cells and focused on a previously undescribed stromal population that we termed Pax7^FAPs^. Mechanistically, we found that Notch signaling biases neonatal Pax7^+^ cells toward a fibrogenic identity at the expense of myogenesis. This lineage plasticity undergoes a rapid age dependent restriction, progressively declining during the first 20 postnatal days and becoming essentially undetectable in 8-week-old adult mice, consistent with Pax7^+^ cells behaving as functionally unipotent myogenic stem cells.

## Results

### Short lineage tracing of Pax7 cells during postnatal hindlimb muscle development at single-nucleus resolution

To track the fate and developmental potential of postnatal *Pax7*^+^ cells, we generated compound *Pax7*^*CreERT2*^; *Rosa26*^*HTB::GFP*^ mice, which upon tamoxifen injection enable the genetic labelling of Pax7-expressing cells and their descendants with nuclear GFP-tagged Histone 2B (H2B-GFP) (Figures 1A and S1A, B). For our transcriptomic profiling, we opted for nuclear instead of cellular isolation due to our previous findings demonstrating significant transcriptional modifications induced during the cell dissociation procedure ^21-23^. Furthermore, single-cell preparations do not capture the nuclei of the multinucleated myofibers, which are a prominent cell type in the myogenic lineage. To ensure successful isolation of labelled nuclei, we utilized H2B-GFP as it provides greater stability after nucleus extraction compared to free, nuclear fluorescence markers (Figure S1A) ^24^. For the snRNA-seq analysis, we isolated DAPI^+^GFP^+^ nuclei by fluorescence-activated cell sorting (FACS) at postnatal days (P) P3, P10, and P22, 48 hours after the initial tamoxifen injection. Nuclei from the entire hindlimb muscles at P3 (WP3) were also sequenced for reference. We chose to study this developmental window since P21 has been suggested to mark the end of nuclear accretion for the bulk of the myocytes, although the precise dynamics remain unclear ^5,25,26^. Our experimental setup was validated by the identification of the core myogenic lineage that originates from Pax7 cells and progresses towards differentiated myocytes before fusing to *Ttn*^+^ myonuclei (Figures 1B and S1C-F). Unexpectedly, in addition to the myogenic lineage, we uncovered cell populations of non-myogenic identity in the descendants of postnatal *Pax7*^+^ cells (Figures 1C, D and S1C-F). Notably, we observed prominent clusters of FAPs (*Dcn*^*+*^), tenocytes (*Tnmd*^*+*^), cells with dual identity (MyoFAPs), and endothelial cells (*Flt1*^*+*^), which progressively decreased with age (Figures 1C and S1C, E). Overall, these data suggest the descendants of Pax7 cells during postnatal muscle growth are mostly, but not entirely, of myogenic identity.

**Figure 1.**
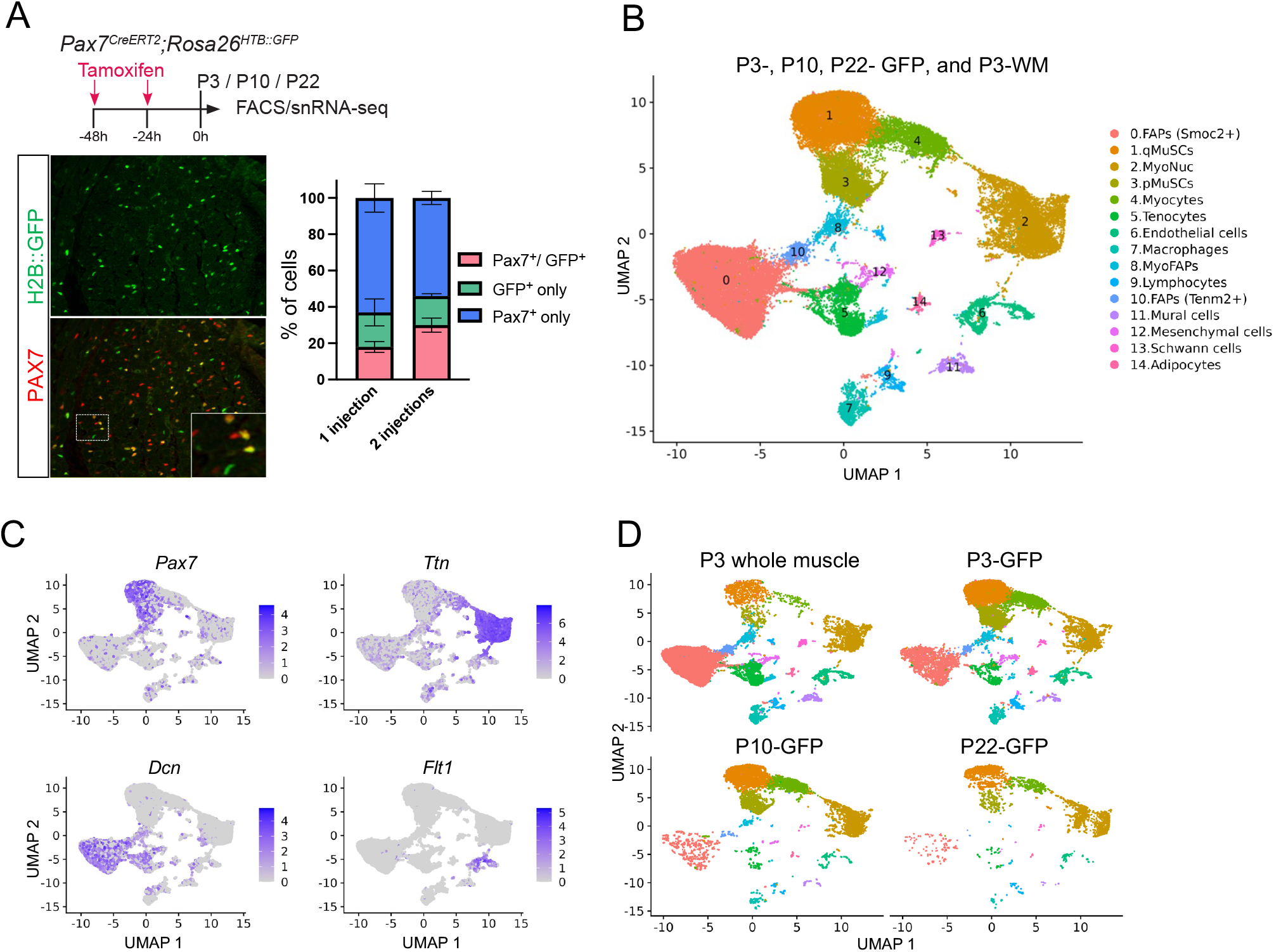
Lineage tracing and single-nucleus transcriptomic profiling of Pax7^+^ cells during postnatal hindlimb muscle development. **(A)** Experimental scheme for labeling the Pax7 cells and their progeny using *Pax7*^*CreERT2*^*;Rosa26*^*HTB::GFP*^ mice. Sections of *Tibialis Anterior* muscle from P3 mice injected twice stained for PAX7 and GFP. *Graph*: labeling efficiency of Pax7 stem/progenitor cells and their progeny in *Pax7*^*CreERT2*^; *Rosa26*^*HTB::GFP*^ pups that received either one tamoxifen injection at-48 h or two injections at - 48 h and-24 h before collection at P13. Pax7+GFP+, Pax7+ only, and GFP+ only cells were quantified ±SD (n=3 mice). **(B)** Integrated UMAP (Uniform Manifold Approximation and Projection) of snRNA-seq experiments, including Pax7-lineage cells from hindlimb muscles collected at P3, P10, and P22 (48 h chase), as well as whole hindlimb muscles from P3 neonatal mice. **(C)** Feature plots displaying the expression profiles of *Pax7, Ttn* (myonuclei), *Dcn (*FAPs), and *Flt1* (ECs) genes in the integrated UMAP. **(D)** UMAP visualizations for each time point.

### Molecular signature of MuSCs and Pax7-derived FAPs across postnatal development

During postnatal development, muscle increases in size by the fusion of differentiated myoblasts onto existing myofibers, while a fraction of PAX7^+^ stem/progenitor cells continues to proliferate to generate more myoblasts and MuSCs ^26^. Higher-definition reclustering of the proliferating MuSCs population (cluster 3) from the integrated UMAP shown in figure 1B, clearly segregated P3 cells (both P3-GFP and WP3) from later stages, whereas P10 cells clustered with P22 cells (Figures 2A, B). Expression of Pax7 and the proliferation marker *Mki67* was comparable across stages; however, P3 cells expressed higher levels of FAP-associated genes, such as collagen type I (*Col1a1*) and tenascin C (*Tnc*), as well as the adipogenic regulator *Pparg* (peroxisome proliferator-activated receptor gamma) (Figure 2C). The distinct molecular profile of early neonatal cells compared to older ones aligns with the observed age-related decline in their capacity to generate alternative lineages.

**Figure 2.**
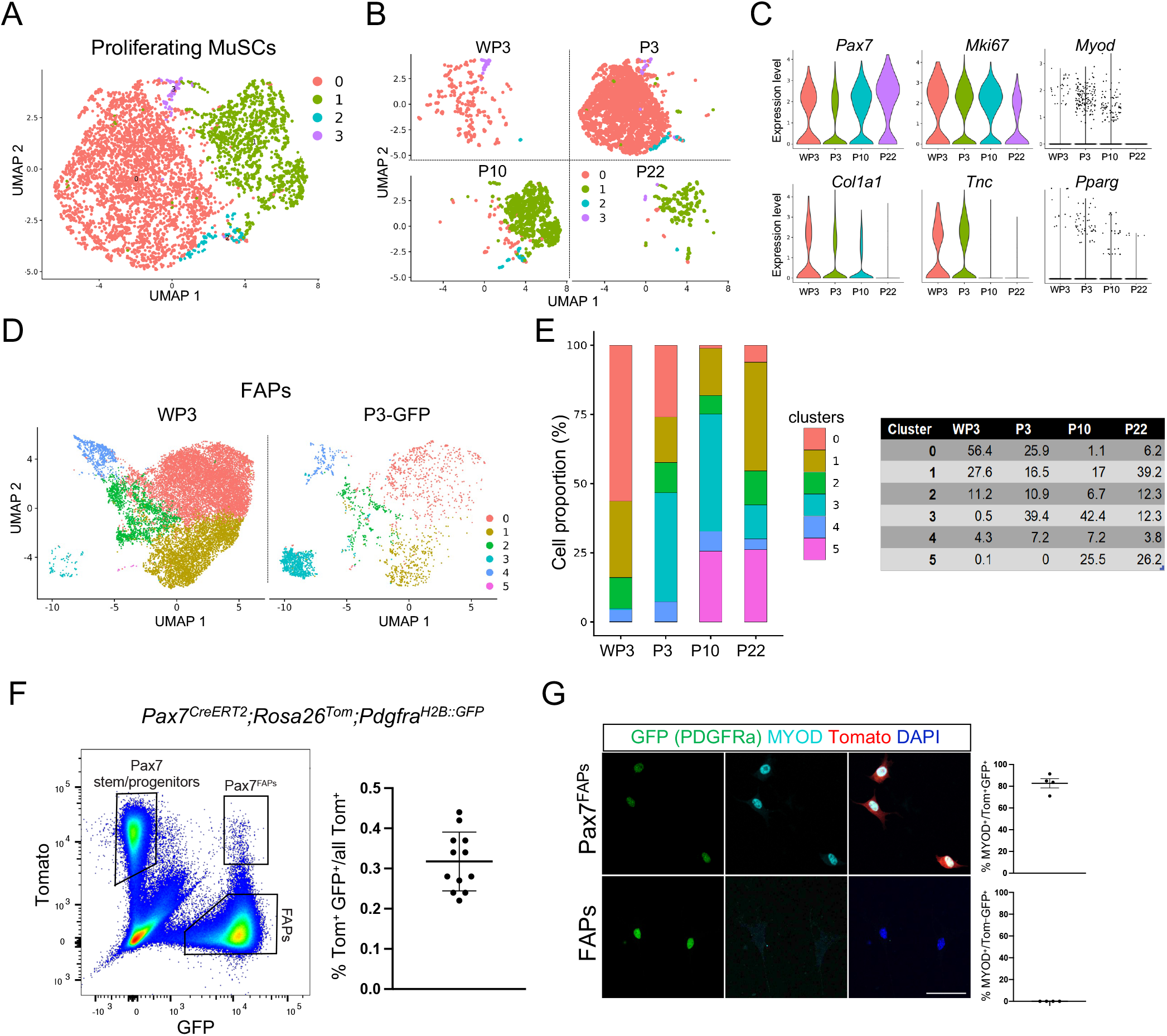
Characterization of proliferating MuSCs and Pax7-derived FAPs postnatal muscle. **(A)** Higher definition clustering of the proliferating MuSCs population extracted from the integrated UMAP (Figure 1B). **(B)** UMAP representations of individual samples. **(C)** Violin plots depicting gene expression of myogenic and proliferation markers (*Pax7, Mki67, Myod1*) and non-myogenic cell markers (*Col1a1, Tnc, Pparg*) across time points. **(D)** Higher definition clustering of FAPs clusters for whole muscle (WP3) and P3-GFP conditions, extracted from figure 1B. **(E)** *Left*: Bar graph showing the proportion of each cluster per time point. *Right*: Table displaying the absolute cell numbers in each cluster. **(F)** FACS analysis on the hindlimb muscles of *Pax7*^*CreERT2*^*;Rosa26*^*Tom/+*^*;Pdgfra*^*H2B::GFP/+*^ mice induced at P1 and P2 and collected at P3. FACS was based on Tomato (Pax7 lineage) and GFP (*Pdgfra* gene) expression. Graph shows the percentage of double positive Pax7^FAPs^ (GFP+ cells within the TOM+ population), (n=12 mice). **(G)** Isolation and culture of the Pax7^FAPs^ (Tom+GFP+) and *bona fide* FAPs (Tom–GFP+) populations shown in (F). Cells were cultured for 6 days and stained for the myogenic factor MYOD and for GFP (*Pdgfra* reporter), (n=4 mice). Percentage (%) of MYOD+ cells is presented over total Tom+GFP+ cells and Tom–GFP+ cells (Pax7^FAPs^ and *bona fide* FAPs, respectively). Data are mean ± SD; scale bar 20 µm.

Despite rapid postnatal tissue growth, a distinct cluster of quiescent MuSCs (qMuSCs) is already identifiable by P3. For the comparison of quiescent cells, adult MuSCs from resting *Tibialis anterior* muscles were integrated in the dataset and used as reference ^22^ (Figure S2A). Higher-definition reclustering identified distinct states of quiescence and separated cells according to developmental stage, with a progressive transition of early neonatal cells towards the adult MuSCs (Figures S2B, C). This maturation is characterized by a shift in gene expression: canonical adult markers such as *Pax7, Chodl*, and *Bmp6* increase gradually with age. In contrast, qMuSCs from younger animals are defined by enriched collagen gene expression and higher levels of the cell cycle inhibitor p57 (*Cdkn1c*) (Figure S2D). This developmental continuum is further supported by “Perinatal” and “Adult” quiescence module scores. These modules (calculated as the median expression of stage-specific marker genes, Table S1) capture the shifting molecular signature of quiescence from P3 through adulthood (Figures S2E, F).

In addition to the expected muscle lineage, we identified non-myogenic cells derived from Pax7 progenitors. The identification of a prominent stromal lineage, which we termed Pax7^FAPs^, raised the question of whether these cells are distinct from *bona fide* FAPs residing in muscle. To address this, we re-analyzed the FAP compartments from the integrated UMAP (Figure 1B) and resolved six subclusters (0-5) (Figures 2D and S2G, H). Pax7^FAPs^ were not evenly distributed across these populations but instead showed a strong enrichment in a single subset (cluster 3) (Figures 2D, E). This cluster was defined by the combined expression of basement membrane-associated collagens (*Col15a1* and *Col4a1*) together with markers typically associated with muscle stem cells, including *Col19a1* and *Pax7*. Importantly, *Col15a1* and *Col4a1*/2 were also expressed in other FAP subpopulations (clusters 4 and 5), indicating that basement membrane gene expression alone is not sufficient to distinguish Pax7^FAPs^ from the broader FAP landscape. Rather, the co-expression of MuSC-associated markers was unique to the Pax7^FAP^-enriched subset. *Col15a1*^+^ mesenchymal cells have already been reported in P0 neonate muscle ^27^ and also as a “universal” subpopulation across tissues, positioned hierarchically between upstream *Dpp4*^+^/*Pi16*^+^ progenitors and more specialized mesenchymal states ^28^. This raises the possibility that the Pax7^FAP^-enriched cluster relates to this broadly conserved intermediate mesenchymal cells, while acquiring a Pax7/MuSC-like signature specific to muscle. Notably, the Pax7^FAPs^ were clearly separable from Myo-FAPs, which formed an independent cluster in the global analysis (Figure 1B) and were therefore excluded from this stage-resolved FAP reclustering.

### Isolation of the Pax7^FAP^ population

To further analyze Pax7^FAPs^ in postnatal hindlimbs, we used a series of additional assays and mouse models. First, we assessed PDGFRa protein expression in Pax7-traced progeny by flow cytometry. To mark the Pax7 lineage, neonatal *Pax7*^*CreERT2*^; *Rosa26*^*HTB::GFP*^ mice were induced with tamoxifen at P1 and P2, hindlimb muscles collected 5 days later, and the dissociated cells were stained with anti-PDGFRa antibody. PDGFRa^+^ cells were found to comprise on average 5.3% of the total GFP^+^ population, consistent with our snRNA-seq-based data (Figure S2I). In addition, to mark and isolate the Pax7^FAPs^, we generated a compound mouse harboring a nuclear GFP reporter of *Pdgfra* gene activity (*Pdgfra*^*H2B::GFP*^), together with alleles to conditionally mark the Pax7^+^ cells and their progeny with the Tomato fluorescent protein (*Pax7*^*CreERT2*^; *Rosa26*^*Tom*^; *Pdgfra*^*H2B::GFP*^). Hindlimb muscles were processed, and cells were sorted based on GFP and Tomato (Tom) expression. A Tom^+^GFP^+^ population corresponding to the Pax7^FAPs^ was readily identified (Figures 2F and S2J). The cells were cultured and stained for GFP (*Pdgfra* reporter), Tom (*Pax7* lineage), and the myogenic marker MYOD. After 6 days in growth medium, FAPs (Tom^−^GFP^+^) retained high GFP expression and were negative for MYOD. Instead, the majority of the Pax7^FAPs^ (Tom^+^GFP^+^) turned MYOD on (81%±SD%), likely reflecting the plasticity of an upstream bipotent progenitor (Figure 2G).

### Implication of Notch signalling in cell priming toward a fibrogenic molecular identity

A key question emerging from these findings was which signals govern fate specification in neonatal Pax7-derived cells. We therefore investigated a potential role for Notch signaling, a well-established regulator of cell fate decisions and a pathway with documented functions in muscle stem cells ^29-31^. To test Notch involvement in our system, we generated *Pax7*^*CreERT2*^; *Rosa26*^*stop-NICD-nGFP*^ mice to drive expression of constitutively active Notch1 (Notch intracellular domain; NICD) in Pax7 stem/progenitor cells and compared them with littermate controls (*Pax7*^*CreERT2*^; *Rosa26*^*stop-dtTom*^). Tamoxifen was administered from P1 to P3 (once every 24 sh) and cells were isolated by FACS at P7 based on fluorescent reporter expression. Purified cells were then subjected to single-cell RNA-seq (scRNA-seq) (Figure 3A). Here, we used single-cell rather than single-nucleus RNA-seq as in the previous experiments, because the EGFP signal of the NICD-ires-nEGFP allele is not preserved during nuclei extraction, in contrast to H2B-GFP reporters that we used before, whose chromatin-tethered fluorescence remains highly stable. Notch activation was confirmed by induction of canonical targets *Hes1* and *HeyL* (Figure 3B) ^31^.

**Figure 3.**
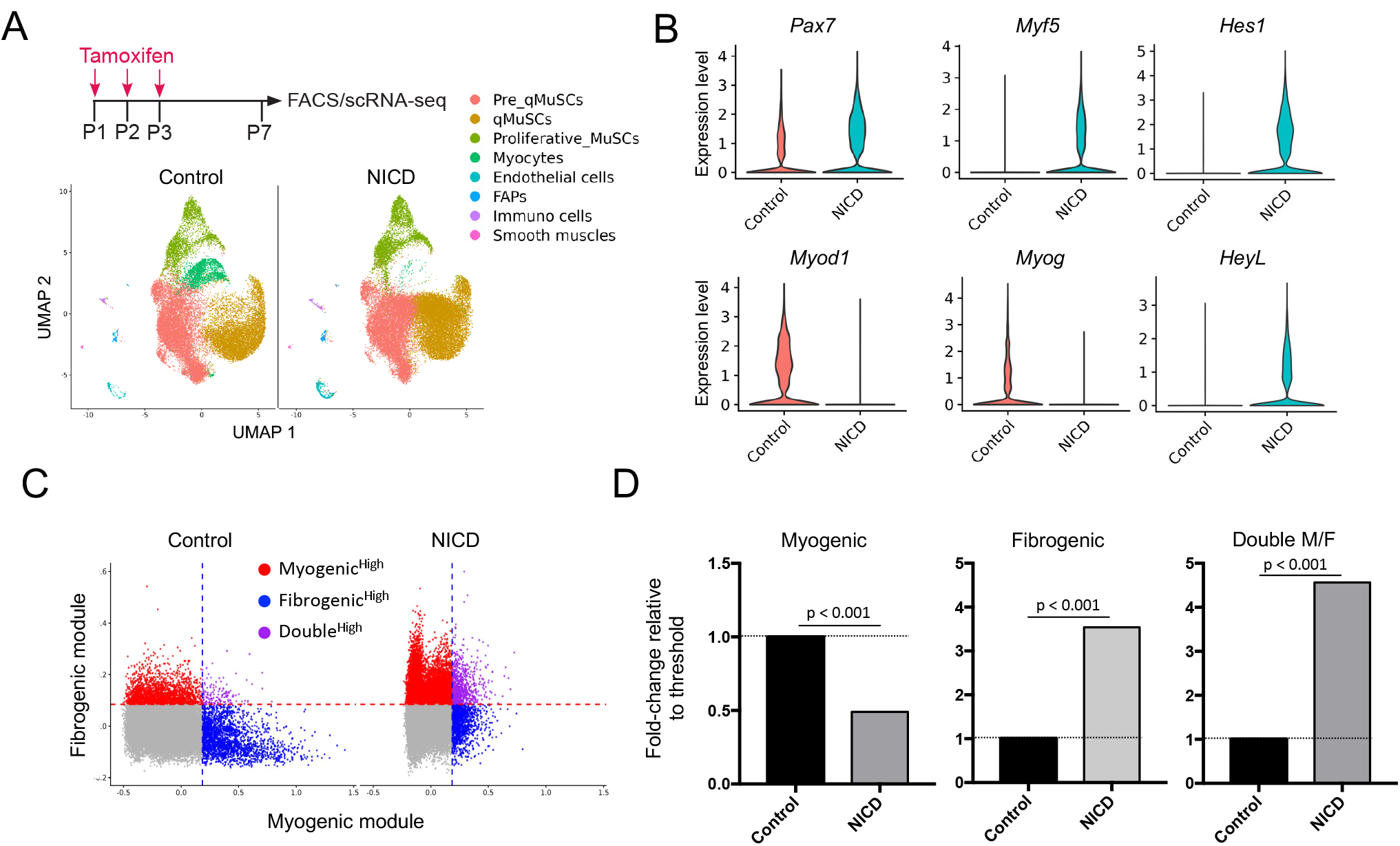
Notch activation primes postnatal Pax7 cells towards a rogenic fate at the expense of myogenesis. **(A)** Experimental scheme: tamoxifen was administered by intraperitoneal injections to *Pax7*^*CreERT2*^*;Rosa26*^*dtTom*^ control mice and *Pax7*^*CreERT2*^*;Rosa26*^*NICD-nGFP*^ mice at postnatal days 1 to 3. Muscles in hindlimbs were harvested at postnatal day 7 (NICD: n = 8 pups; Control: n = 6 pups). Integrated UMAP of single-cell RNA sequencing from control and NICD samples. **(B)** Violin plots showing expression levels of myogenic stem cell (*Pax7, Myf5*) differentiation (*Myod1*, and *Myog*), and Notch signaling activity reporter (*Hes1, HeyL*) genes across all cells in control and NICD datasets. **(C)** Scatter plot of myogenic and fibrogenic module scores computed as the median expression of genes using AddModuleScore. Fibrogenic score: *Pdgfra, Dpp4, Smoc2, Fap, Ebf1, Ebf2, Ly6a, Dcn, Cd34, Mmp14, Col3a1, Col1a1, Col1a2, Fn1, Lox, Postn, Pi16, Ly6c1, Col15a1, Penk, Ccl19, Ccl21a, Bst1, Cxcl12, Lepr, Sp7, Npnt, Ces1d, Comp, Fmod, Coch, Wt1, Fbln1, Sfrp1, Hhip, Aspn*, and *Bmp4; M*yogenic score: *Myod1, Myf5, Myog, Myh3, Myh8, Mymk*, and *Mymx*. Cells with scores above the top 10% threshold in Control are defined as high-score cells. **(D)** Bar graph comparing the population of high-score cells, normalized to control (set to 1). Since the biological replicates were pooled and sequenced together, each condition was treated as a pseudo-bulk population and compared the proportion of cells exceeding the Control-defined top 10% threshold using Fisher’s exact test. For all scores, statistical comparisons between Control and NICD showed significant differences (P<0.001).

Our previous work showed that Notch activation exerts context-dependent effects in muscle progenitors: it blocks myogenic commitment and expands embryonic progenitors (E12.5), whereas in fetal progenitors (E18.5) it promotes a cell-cycle exit ^32^. Consistently, NICD overexpression in postnatal Pax7^+^ cells produced a phenotype resembling the fetal setting with respect to the myogenic lineage. We observed a marked reduction in the terminally differentiated Myogenin^+^ population, accompanied by a strong expansion of the quiescent MuSC compartment (Figure 3A, B). Of interest, NICD induction in the neonatal context induced a mesenchymal/FAP-like transcriptional signature within Pax7^+^ cell population (Figures 3C, D and S3A, B). While overall FAP production was only modestly increased (Figure S3A), Pax7^+^ cells overexpressing NICD were notably enriched for mesenchymal and fibroblast-associated markers (Figures 3C, D). Notably, sustained Notch activation also enriched for cells with a double fibrogenic/myogenic identity (Figures 3C, D). Together, these data indicate that Notch signaling primes neonatal Pax7^+^ cells toward a fibrogenic molecular identity at the expense of myogenic differentiation, thereby biasing their trajectory toward the Pax7^FAPs^ lineage.

### Pax7^FAPs^ in the adult skeletal muscles

We then asked if the Pax7^FAPs^ generated at early postnatal stages represent a transient developmental state or instead a stromal cell population persisting into adulthood. To address this, we labelled Pax7^+^ cells during early postnatal stages (P1 to P4) in *Pax7*^*CreERT2*^; *Rosa26*^*HTB::GFP*^ mice and analyzed their progeny 2 months later (Figure 4A). First, GFP^+^ lineage-traced cells were isolated in adult mice by flow cytometry, using established combinations of antibodies for endothelial cells, FAPs and MuSCs (Figure 4A) ^33^. As shown in figures 4B and 4C, we could readily identify GFP-marked MuSCs (96%±SD% of GFP^+^ cells), FAPs (0.35%±SD% of GFP^+^ cells), and endothelial cells (0.51%±SD% of GFP^+^ cells). Moreover, by immunofluorescence on TA muscle sections, we could detect GFP^+^ cells residing inside vessels, marked by CD31 expression, and also in the interstitial space (Figures 4D and S4A). To further characterize the long-lineage descendants in the adult muscle, we performed snRNA-seq on FACS-isolated GFP^+^ nuclei from hindlimb muscles (Figures 4E and S4B). As expected, most of the isolated nuclei were myofiber nuclei (89.6% of total nuclei), including neuromuscular junction (NMJ) and myotendinous junction (MTJ) myonuclei, whereas a separate population of quiescent MuSCs was isolated. Notably, non-myogenic cells of FAP, immune and endothelial profile were also identified (Figure 4E). Comparison of the Pax7^FAPs^ with the *bona fide* FAPs showed no enrichment towards a specific subpopulation. Therefore, the Pax7^FAPs^ generated at early postnatal stages establish residence within adult muscle, albeit in limited quantities and without any particular transcriptional signature.

**Figure 4.**
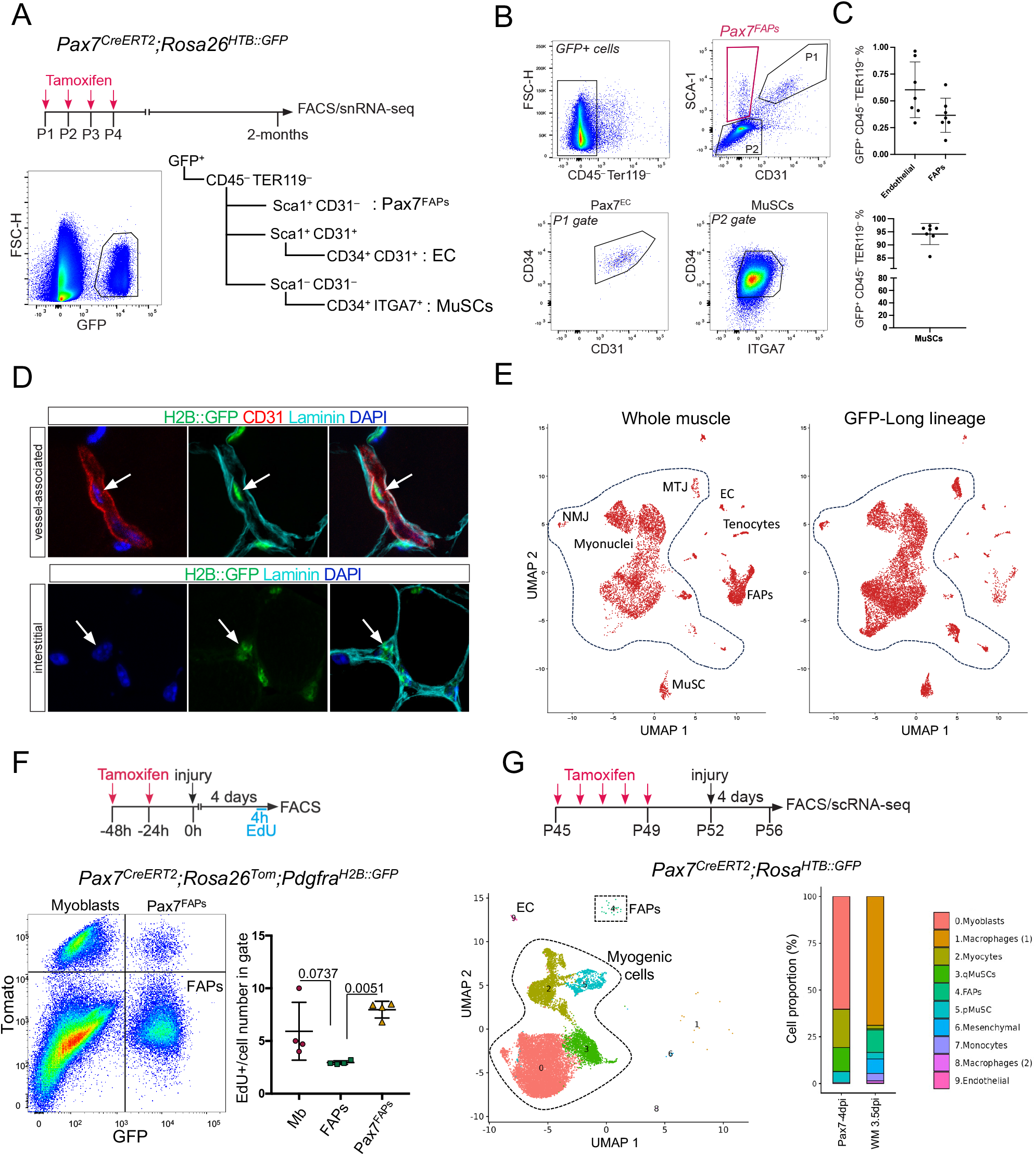
Adult Pax7^FAPs^ persist into adulthood and emerge after injury. **(A)** Experimental strategy for 2-month lineage tracing of Pax7 cells; *Pax7*^*CreERT2*^; *Rosa26*^*HTB::GFP*^ mice were induced with tamoxifen at P1-P4 and sacrificed at 2 months of age. Different muscle resident cell populations were analyzed by FACS. Explanatory tree of the antibody panel used: the GFP+ cells (left), negative for the hematopoietic markers CD45, TER119, were further gated to identify endothelial cells (Sca-1+CD31+CD34+), FAPs (Sca1+CD31–and MuSCs (Sca-1–CD31–CD34+ ITGA7+). **(B)** FACS profiles of the GFP+ cells shown in (A). The Pax7-derived FAPs are highlighted with a red gate. **(C)** Percentage of GFP+ cells in the endothelial, FAP, and MuSC populations; values are mean ± SD (n=7 mice). **(D)** Representative IHC pictures from adult *Pax7*^*CreERT2*^*;Rosa26*^*HTB::GFP*^ mice induced with tamoxifen at P1 to P4. Immunostaining was performed on TA sections for GFP (Pax7 lineage) and Laminin (2 months old mice) or for GFP (Pax7 lineage), Laminin and CD31 (5.5 weeks old mice) to identify the presence of interstitial and endothelial-associated cells, respectively. Arrows point to Pax7 lineage-traced GFP+ cells. **(E)** snRNA-seq after long lineage tracing *in vivo. Pax7*^*CreERT2*^*;Rosa26*^*HTB::GFP*^ mice were induced at P1-P4 and sacrificed at 2-months-old. SnRNA-seq was performed on DAPI+GFP+ nuclei from hindlimb muscles (n=4 mice pooled). **(F)** Experimental strategy to compare bona fide FAPs and Pax7^FAPs^ in response to injury. Pax7^CreERT2^; Rosa26^Tom/+;^ Pdgfra^H2B::GFP/+^ mice (2-2.5 months old) were induced 48 hours prior to muscle injury, and injury was applied to the gastrocnemius and TA muscles. EdU was administered intraperitoneally 4 hours before sacrifice. Graph: The proportion of EdU-positive cells was calculated among Myoblasts (Mb) (Tom^+^GFP^−^), Pax7^FAPs^ (Tom^+^GFP^+^), and bona fide FAPs (Tom^−^GFP^+^) based on absolute cell numbers. Quantification was performed using absolute cell counts obtained by FACS analysis (n = 4 mice). **(G)** Lineage tracing of Pax7+ cells 4 days post injury by single-cell RNAseq in adult mice. Pax7 cells were genetically marked in resting muscle with H2B::GFP and labeled cells were isolated and sequenced 4 days after BaCl_2_-induced injury. 99.2 % of cells consisted of myogenic cells, of 0.6 % endothelial cells, and of 0.1% FAPs.

Previous reports have shown that a small fraction of adult MuSCs aberrantly acquire a fibrogenic fate during regeneration, and that this fraction increases markedly in *mdx* dystrophic mice and in aged (18-month-old) mice ^15,16^. To trace these cells and test their potential involvement in muscle regeneration, adult *Pax7*^*CreERT2*^; *Rosa26*^*Tom*^; *Pdgfra*^*H2B::GFP*^ mice were induced with tamoxifen and injured the TA and gastrocnemius muscles with BaCl_2_ 24 h after the last tamoxifen injection. Muscles were collected 4 days after injury, with EdU administered 4 h before collection to label cycling cells. Pax7^FAPs^ were identified by FACS as Tom^+^GFP^+^ cells and compared with FAPs (Tom^−^GFP^+^) (Figures 4F and S4C). Quantification of EdU showed that the Pax7^FAPs^ were actively cycling and, notably, proliferated more than *bona fide* FAPs (Figure 4F).

These results raised the question of whether adult Pax7-expressing cells retain the multipotency observed at neonatal stages or instead become restricted to the myogenic lineage. To address this more broadly, we lineage-labeled Pax7^+^ cells in resting *Pax7*^*CreERT2*^; *Rosa26*^*HTB::*GFP^ mice and induced injury in the gastrocnemius and TA muscles with BaCl_2_. Four days after injury, lineage-traced GFP^+^ cells were isolated and subjected to single-cell RNA sequencing (Figure 4G). As expected, the vast majority of GFP^+^ cells were myogenic, including MuSCs, myoblasts, and myocytes (myonuclei are not captured by single-cell analysis). A small fraction of non-myogenic cells was also detected, including rare FAPs (consistent with previous reports and with our data), as well as few endothelial cells (Figures 4G and S4D). Together, these results indicate that, in adult muscle the differentiation potential of Pax7^+^ cells is largely restricted to myogenesis.

## Discussion

The experiments of this study focused on hindlimb muscles during postnatal development. In the limb, embryonic myogenesis occurs between E10.5 and 12.5 in mouse and begins when Pax3^+^ muscle progenitors delaminate from the hypaxial dermomyotome of somites and migrate into the limbs (E10.5) ^4^. The migrating Pax3^+^ population will initiate the myogenic program with expression of *Myf5* and *Pax7* (expression at E11.5), will proliferate and eventually form limb muscles ^34,35^. In this study, we demonstrate that postnatal *Pax7*^*+*^ cells that did not arise in the dermomyotome but in the limb, can give rise to non-myogenic populations, among them populations with a FAP signature (Pax7^FAPs^).

FAPs belong ontologically to the general category of muscle connective tissue (MCT) fibroblasts and they differ in their developmental origin, depending on their anatomical location ^36^. Experiments in the mouse and the chick show that limb MCT fibroblasts derive from the lateral plate mesoderm (LPM) ^37,38^ and during limb formation, LPM-muscle connective tissue acts to ensure correct muscle patterning ^39-41^. The notion that LPM-derived cells in the limb give rise strictly to MCT, whereas myogenic populations are of somitic origin, was recently challenged by studies that showed a localized LPM-derived cell contribution to muscle tissue, close to the MTJ ^8,9^. Single-cell RNA-seq at P0 at the MTJ area identified a population with a dual fibrogenic/myogenic molecular identity, including MTJ and tenocyte markers. In our postnatal snRNA-seq datasets, we identified Pax7-derived FAPs and tenocytes, that form distinct clusters from the myogenic ones. We believe that we target a fraction of the cells described by Esteves de Lima et al. (8) and Yaseen et al. (9), since the dual identity population described in these studies, expresses *Pax7*. However, the Pax7-derived FAPs and tenocytes that we identified have distinct properties since they persist into adulthood.

We identified additional non-myogenic populations in our dataset such as adipocytes, mural cells, macrophages and Schwann cells. The presence of adipocytes may indicate that Pax7-derived FAPs can undergo terminal differentiation similarly to *bona fide* FAPs ^42^, while a Pax3^+^ precursor and/or a DM origin for endothelial and mural cells has previously been described, showing that muscle/stem progenitors can give rise to these populations ^34,43-45^. Moreover, since some bone marrow progenitors have been shown to be capable of myogenic potential ^46^, we can speculate that the opposite is happening for the macrophages that we observe. However, we believe that the Schwann cells that we identify here, due to their developmental origin and their small size in our dataset, are likely the result of background signal. The combination of a strong reporter line (H2B::GFP) with single-nucleus RNA-seq likely places the analysis close to the sensitivity threshold, resulting in low-level transcriptional noise.

Several studies have described mesenchymal cells heterogeneity within individual tissues. In the skeletal muscle, FAP heterogeneity at the transcription level has also been reported during homeostasis, regeneration, and in dystrophic context ^28,47-54^. Our analysis showed a significant enrichment of Pax7^FAPs^ towards a cluster marked by *Col15a1*^+^ and *Col4a1/2*^+^, together with markers associated with MuSCs, like *Col19a1* and *Pax7*. This subpopulation was previously identified as a universal subpopulation in a study that compared mesenchymal cells across 17 murine tissues ^28^. RNA trajectory analysis proposed a model where mesenchymal cells initiated from the *Pi16*^*+*^*/Dpp4*^*+*^ population, passed through the *Col15a1*^*+*^ population, and gave rise to more specialized cells, indicating that they are developmentally linked. According to this trajectory, the *Pi16*^*+*^ population is placed in the highest hierarchical position and the *Col15a1*^+^ population in the second highest, in agreement with the *Dpp4*^*+*^ upstream mesenchymal population described previously ^49^. Our data, however, suggest that in the skeletal muscle Pax7^+^ cells is an alternative source of the *Col15a1*^*+*^ population.

Pax7-derived FAPs, endothelial and immune cells in the limb could support postnatal development and autonomously protect MuSCs. We cannot rule out the possibility that MuSCs may partially exhibit transient plasticity in response to the highly dynamic postnatal environment. Pessina et al. (2015) showed that MuSCs can acquire a fibrogenic identity in the TGFb-rich microenvironment of *mdx* dystrophic mice. Moreover, a switch of MuSCs to a fibrogenic identity in aged mice was previously reported ^55^. Signs of plasticity in the myogenic lineage are also manifested during the regeneration of healthy tissue, as shown by the presence of ‘immunomyoblasts’ co-expressing myogenic and immune markers ^51^. It could, therefore, be suggested that in a highly dynamic environment, such as that characterizing postnatal development, MuSCs transiently acquire features towards other fates. However, the fact that some of the populations that we identify persist into adulthood and Pax7^FAPs^ specifically demonstrate a remarkable expansion upon muscle injury compared to *bona fide* FAPs, suggests otherwise.

Interestingly, the reverse phenomenon, that is pervasiveness of other types of mesodermal cells into developing skeletal muscle, has been previously described. As such, embryonic cells of the dorsal aorta with vascular cell characteristics, named meso-angioblasts, have been shown to contribute to skeletal muscle growth ^56^ and, following transplantation into chick embryos, to incorporate mesodermal tissues such as cartilage, bone, smooth muscles ^57^. Likewise, we have previously demonstrated that fetal endothelial cells residing within the developing skeletal muscles can differentiate into myotubes *in vitro* and fuse with adult myofibers following transplantation ^58^. Additionally, interstitial cells that express *Twist2, Hoxa11*, or both, which can fuse with myofibers and contribute to muscle growth, have been described ^59,60^.

We propose that postnatal skeletal muscle stem/progenitor cells possess a certain degree of phenotypic plasticity. By combining *in vivo* lineage tracing with targeted single-cell technologies, our study reveals that Pax7-expressing stem/progenitor cells, during postnatal limb muscle development, are not restricted to a single lineage as previously thought. Instead, they demonstrate multipotency, differentiating into various lineages. The reasons behind the multipotent identity of postnatal Pax7^+^ cells in muscle and whether there is functional difference among Pax7-derived cells compared to their homologous populations, remain unknown. Nonetheless, our findings indicate that these Pax7-derived non-myogenic cells contribute to the cellular architecture of muscle from early postnatal stages through adulthood. Altogether, this study advances our understanding of tissue composition and development during postnatal muscle growth.

## Figure legends after each corresponding figure

## ACKNOWLEDGEMENTS

The experiments in the Mourikis and Relaix teams were supported by funding from the Agence National de la Recherche (ANR) ANR-19-CE14-0008, ANR-22-CE13-0023-02 and ANR-22-CE13-0023-02, the Association Française contre les Myopathies (AFM) via Translamuscle (project 22946), LabexREVIVE (ANR-10-LABX-73), and the PhD fellowships from the Fondation pour la Recherche Médicale (FRM) and the Université franco-allemande (UFA). T.H. was supported by the Uehara Memorial Foundation (202340029).

The research conducted in the Le Grand lab was supported by funding from the Inserm, the CNRS, the Agence National pour la Recherche (ANR-19-CE13-0016), the AFM-Téléthon (MyoNeurAlp2 project) and the European Joint Program for Rare Diseases (EJPRD JTC2019, Myocity project).

We are grateful to Dr Glenda Comai for her scientific support during the project. We thank the IMRB Cytometry platform, and the Genom’IC facility (Cochin Institute, Paris) and the Imagine Single-Cell platform (Imagine Institute, Paris) for the library preparation and sequencing.

## AUTHOR CONTRIBUTIONS

S.G., T.H., P.M, and F.L.G designed and interpreted the experiments. P.M., F.R. and F.L.G. obtained funding. M.D.S. and S.J. performed the P22 snRNA-seq experiments, together with S.G. and T.H.; T.H., S.F. and F.L.G. performed the bioinformatics analyses. All authors reviewed and edited the manuscript.

## METHODS

### RESSOURCE AVAILABILITY

#### Lead contact

Further information and requests for resources and reagents should be directed and will be fulfilled by the lead contact, Philippos Mourikis.

#### Materials availability

This study did not generate new unique reagents. Nonidet™ P40 Substitute (Sigma-Aldrich, 74385) used in this study has been discontinued. The same applies to the 96-well plates (Neobits, 10204961).

#### Data and code availability

Microscopy data reported in this paper will be shared by the lead contact upon request. This paper does not report original code. The data discussed in this publication have been deposited in NCBI’s Gene Expression Omnibus and are accessible through GEO Series accession number GSE280693. Any additional information required to reanalyze the data reported in this paper is available from the lead contact upon request.

### EXPERIMENTAL MODEL AND SUBJECT DETAILS

#### Mouse strains

The following mouse lines were kindly provided by the corresponding laboratories as described: *Tg:Pax7-nGFP* (S. Tajbakhsh ^61^, The Jackson laboratory, stock 036759), *Pax7*^*CreERT2(Gaka)*^ (G. Kardon ^14^, The Jackson laboratory, stock 017763), *Tg:Pax7-CreERT2* (S. Tajbakhsh; ^30^), *Rosa26*^*stop-NICD-nGFP*^ (D. Melton ^62^, The Jackson laboratory, stock 008159), and the reporter lines *Pdgfra*^*H2B::GFP*^ (S. Tajbkahsh ^63^, The Jackson laboratory, stock 007669), _*Rosa26*_*Lox-stop-LoxHTB* _(C. Birchmeier_ 64_), *Rosa26*_^*mTomato-stop-mGFP*^ (S. Tajbakhsh ^65^, The Jackson laboratory, stock 007576) and *Rosa26*^*TdTomato*^ (S. Tajbakhsh ^66^). Male and female mice were used for all the studies, except for snRNA-seq at P22 where only female mice were used to match our previous snRNA-seq datasets from Machado et al., 2021 (22) in adult mice. For each experiment presented, age is indicated in the corresponding legend. Animals were handled as per French and European Community guidelines and protocols were approved by the ethics Committee at the French Ministry Project No: 20-027 #24357).

#### Primary cell cultures

Bulk hindlimb muscle cell dissociation was performed according to the protocol of Machado et al., 2017 (21) for non-fixed tissues. For experiments performed at P3, muscle bulk filtration upon digestion was done only with 70 µm filters (Miltenyi biotec, 130-110-916) into 15 ml tubes and the last centrifugation was performed at 600 g for 8 min. Cells sorted by FACS were plated on 96 well plates suitable for microscopy (Neobits, 10204961) coated with Matrigel (Corning, 354248) in growth medium with DMEM (GIBCO), 20% fetal bovine serum (FBS), 1% penicillin/streptomycin (GIBCO), and 4 ng/μl basic FGF (bFGF, Peprotech, 450-33). Cells were plated at a density of 80 cells/mm^2^ for MuSCs, and at a density of 9-15 cells/mm^2^ (300-500/well) for Pax7^FAPs^. Bona fide FAPs were plated at a density of 2700 cells/well. Cells were cultured for 6 days at 37°C, 5% CO_2_.

### METHOD DETAILS

#### Tamoxifen administration

For Cre recombination, tamoxifen (Sigma-Aldrich, T5648) was prepared in corn oil/5% ethanol at 37°C in a rotating oven for 1 h, protected from light and at 20 mg/ml stocks. For 48 h short lineage tracing, mice were injected intraperitoneally twice, at 48 h and 24 h before sacrifice (for injections at P1 and P2, administration was intragastric), at 200 µg/g. The same protocol was used for experiment at Figure 3A (7 days chase). All injections for short lineage tracing experiments including snRNA-seq experiments were performed in the morning. For longer cell lineage tracing experiments, a protocol described by Lizen et al., 2015 ^67^ was applied. Specifically, mice were injected intragastrically from P1 to P4 (P1: 50 µg/g, P2: 75 µg/g, P3: 75 µg/g, P4: 100 µg/g) at a final volume of 10 µl each time. Intragastric and intraperitoneal injections were performed every 24 h in late afternoon (19:00-20:00), as it was important, especially in P1, for the mice to have reached a weight of 1.7-1.8 g prior to the first injection. Intragastric injections were facilitated by the transparent skin, which makes the white stomach visible at these stages due to the presence of milk. For all injections, a 100 µl Hamilton syringe was used with customized needle (point type 2, gauge 22s, length 51 mm, ext.: 0.72 mm). To ensure the survival of the pups, it was vital that the gloved hands were embedded in the woodchip to cover the smell of the gloves to avoid cannibalization. In adult mice, tamoxifen was administered at a dose of 150 µg/g of mouse by intraperitoneal injection for five consecutive days, every 24 h.

#### Muscle injury

For muscle injury, mice were anesthetized with administration of 5% isoflurane initially in chambers and then via intranasal administration of 2% isoflurane during the injury. Injury of TA and gastrocnemius muscles was induced by injection of 30 and 40 µl, respectively, of 0.6 % BaCl_2_ (Sigma-Aldrich, 202738)

#### Extraction of single nuclei

For all snRNA-seq experiments, nuclei were extracted from hindlimb muscles according to the protocol we described in detail in Dos Santos et al., 2021. Prior to nuclei loading on the 10xGenomics chip for encapsulation, nuclei were counted using counting chambers Malassez and their quality was checked under the microscope in brightfield according to 10x guidelines. For snRNA-seq at postnatal stages and adulthood, mice were pooled together: at P3 upon lineage tracing, 4 females and 5 males were pooled, at P3, whole muscle, 3 males and 3 females were pooled, at P10 upon lineage tracing one female and one male were pooled, at P22 upon lineage tracing, 8 females were pooled, at adult stage upon long lineage tracing two females and two males were pooled.

#### Library preparation for snRNA-seq

For all snRNA-seq experiments, library preparation was performed according to 10X Genomics guidelines, following the Chromium Next GEM Single Cell 3ʶ Reagent Kits v3.1 (Dual Index) guide. Detailed information can be found in the 10XGenomics guide mentioned above as well as in Dos Santos et al., 2021. Libraries were generated by the Genom’IC facility (Cochin Institute, Paris) and sequenced on the NextSeq 2000 platform (Illumina) to obtain around 50,000 reads per nuclei.

#### Cell sorting for scRNA-seq

scRNA-seq samples were prepared from *Pax7*^*CreERT2*^*-NICD* and control (*Pax7*^*CreERT2*^*-tdTomato*) mice. Tamoxifen was administered at postnatal days P1 to P3, and hindlimb skeletal muscles were harvested at P7. For each genotype, muscles from multiple pups were pooled prior to processing (NICD: n = 8 pups; Control: n = 6 pups). In a separate experiment, samples from muscle injury experiments using *Pax7*^*CreERT2*^*;Rosa26*^*H2B-GFP*^ mice were collected following tamoxifen administration. Injured muscles were harvested at P56, corresponding to 4 days post-injury (n = 4). Bulk hindlimb muscle dissociation was performed according to the protocol described by Machado et al. (2017) for non-fixed tissues. For postnatal samples, digested muscle suspensions were filtered through 70 µm strainers (Miltenyi Biotec, 130-110-916) into 15 ml tubes. The final centrifugation step during tissue preparation was performed at 600 × g for 8 min at 4°C. For FACS, GFP-positive cells (NICD samples) and tdTomato-positive cells (control samples) were isolated. A total of 120,000 cells per sample were sorted. After sorting, cells were centrifuged at 900 × g for 10 min at 4°C prior to downstream scRNA-seq processing.

#### Library preparation for scRNA-seq

For scRNA-seq experiments, library preparation was performed according to 10X Genomics guidelines, following the Chromium Next GEM Single Cell 3ʶ Reagent Kits v3.1 (Dual Index) guide. Libraries were generated by Institute Imagine (Paris) and sequenced on the NovaSeq 6000 platform (Illumina) to obtain around 50,000 reads per cell.

#### FACS analysis based on antibodies

Bulk hindlimb muscle cell dissociation was prepared according to the protocol of Machado et al., 2017 for non-fixed tissues. For experiments performed at postnatal stages, muscle bulk filtration upon digestion was done only with 70 µm filters (Miltenyi Biotec, 130-110-916) into 15 ml tubes and the last centrifugation was performed at 600 g for 8 min at 4°C. Muscle bulks were blocked in 2% BSA in PBS for 15 min, centrifuged for 5 min/600 g at 4°C and were incubated on ice for 45 min protected from light, with the following conjugated antibodies diluted in 0.2% BSA in PBS: anti-TER119-PECy7 (3.5 µl, BD Biosciences, 557853) anti-CD45-PECy7 (3.5 µl, BD Biosciences, 552848), anti-Sca-1-PE (3.5 µl; BD Biosciences, 553108), anti-ITGA7-Alexa Fluor 700 (14 µl, R&D Systems, FAB3518N), anti-CD34-BV605 (14 µl, BD Biosciences, 750918) and anti-CD31-APC (2.66 µl, BD Biosciences, Cat # 551262) in a final volume of 200 µl. After incubation, cells were washed with PBS (600 g/ 5 min/ 4°C) and resuspended in 0.2% BSA in PBS. For experiments where cells were stained only for PDGFRα, the conjugated antibody anti-CD140α-APC (2 µl, eBioscence, 17-1401-81) was used. EdU incorporation was detected using the Alexa Fluor 647 Click-iT Plus EdU Flow Cytometry Assay Kit (Invitrogen, C10634), according to the manufacturer’s instructions.

#### Immunostaining on cells and muscle sections

Cells were washed, fixed with 4% formaldehyde for 10 min at RT and washed again three times. All washes were performed with PBS for 5 min each unless otherwise indicated. Cells when then permeabilized and blocked using a solution of 10% goat serum (Sigma-Aldrich, G9023) and 0.25% Triton in PBS for 1 h at RT on a rocker. After one wash, cells were incubated with primary antibodies listed in Key Resources Table, in 5% goat serum and 0.025% Triton in PBS overnight at 4°C on a rocker. Primary antibody mix was washed off three times and cells were incubated with Alexa-conjugated antibodies (1 in 500) and DAPI (1 in 1000) in PBS for 1h at RT. Cells were washed three times and mounted with fluoromount™ (Invitrogen, 00-4958-02). Prior to muscle section for immunohistochemistry (IHC), dissected muscles were fixed in 2% formaldehyde for 2 hours at 4°C, rotating, washed three times (3×10 min) and placed in 15% sucrose/H_2_O overnight at 4°C rotating. The next day muscles were transferred in 30% sucrose/H_2_O at 4°C until precipitation. Muscles were frozen in liquid-nitrogen-cooled isopentane and cut at 7 µm cryosections that could be stored in-80°C until IHC. For IHC, cryosections were equilibrated at RT (20 min) and rehydrated in PBS (10 min). Upon fixation with 4% formaldehyde (10 min), they were washed three times with PBS and were permeabilized with 0.5% Triton in PBS (12 min) followed by three washes. Blocking was done in 10% FBS, 10% Goat serum (Gibco) and 0.25% Tween in PBS for 1.5 h at RT. Following two washes, sections were incubated with primary antibodies listed in key resources table in 0.1% BSA (Jackson Immunoresearch, 001-000-162) in PBS overnight at 4°C. The next day, sections were washed three times and incubated with Alexa-conjugated antibodies and DAPI (1 in 1000) in 0.1% BSA in PBS for 1 h at RT. Sections were then washed three times and mounted with fluoromount™.

### QUANTIFICATION AND STATISTICAL ANALYSIS

#### High-throughput datasets processing

Data demultiplexed in fastq format was checked with fastqc and fastq_screen to control quality sequencing. Sequencing reads were processed on the 10x Genomics Single Cell 3’ v3 platform, utilizing Cell Ranger version 4.0 to align and count data for each sample. A custom reference genome based on mm10 was modified to incorporate the H2BGFP gene (customref-CR5-mm10-2020-A-H2BGFP), with the “Include introns” option activated to facilitate the detection of immature transcripts. For each age group or experimental batch, the matrices were imported into RStudio, where standard quality control procedures were applied using Seurat v.5.4. Ambient RNA and doublets were systematically filtered out using the DropletUtils and scDblFinder packages, respectively.

Original Louvain algorithm with 0.7 resolution was performed to determine clusters. We used Rreduction and RunUMAP to generate reduced dimension graphs. Following clustering and marker identification, cell populations were manually annotated based on established canonical markers. Four distinct data integrations were performed using either Seurat or Harmony, selecting the method that provided the most robust results. (1) postnatal Seurat objects (GFP-traced and whole muscle nuclei at P3 (WP3), and GFP-traced nuclei at P3, P10 and P22). (2) Seurat objects of adult snRNA-seq from intact TA muscle from Machado et al., 2021 (22) integrated with postnatal WP3, and GFP-traced nuclei at P3, P10 and P22. (3) Long-traced GFP nuclei were integrated with adult snRNA-seq from intact TA muscle from Machado et al., 2021 (22). (4) Seurat objects of scRNA-seq from injured TA muscle at 4 dpi integrated with scRNA-seq from injured TA at 3.5 dpi generated by Oprescu et al., 2020 (51). In all integrations, the optimal dimension threshold was determined by a custom method.

#### Statistical analysis

Experiments involving mice were performed with a minimum of three biological replicates and results are presented as the mean ±SD, unless otherwise indicated. One-way ANOVA with Tukey’s test in figure 4F was performed using GraphPad Prism Software, v9.0.

To compare the enrichment of high-scoring cells, a “high-score” threshold was defined as the top 10% of the Control population. Each condition was treated as a pseudo-bulk population, and the proportion of cells exceeding this threshold was compared using Fisher’s exact test in figure 3D.

For the comparison of module scores across multiple groups in figure S2E, a non-parametric approach was employed due to the non-normal distribution of single-cell expression data. The Kruskal-Wallis rank sum test was used to assess global differences among the three groups. The Kruskal-Wallis statistic (*H*) was reported as a measure of the shift in rank distributions between groups. For post-hoc analysis, Dunn’s multiple comparison test was performed with Benjamini-Hochberg (BH) adjustment to identify specific pairs with significant differences. All statistical analyses were conducted in R (v4.5.2) using the FSA and stats packages.

## References

1 Biressi, S. et al. Intrinsic phenotypic diversity of embryonic and fetal myoblasts is revealed by genome-wide gene expression analysis on purified cells. Dev Biol 304, 633–651, doi:10.1016/j.ydbio.2007.01.016 (2007).

2 Ben-Yair, R. & Kalcheim, C. Lineage analysis of the avian dermomyotome sheet reveals the existence of single cells with both dermal and muscle progenitor fates. Development 132, 689–701, doi:10.1242/dev.01617 (2005).

3 Kassar-Duchossoy, L. et al. Pax3/Pax7 mark a novel population of primitive myogenic cells during development. Genes & development 19, 1426–1431, doi:10.1101/gad.345505 (2005).

4 Relaix, F., Rocancourt, D., Mansouri, A. & Buckingham, M. A Pax3/Pax7-dependent population of skeletal muscle progenitor cells. Nature 435, 948–953 (2005).

5 Lepper, C., Conway, S. J. & Fan, C. M. Adult satellite cells and embryonic muscle progenitors have distinct genetic requirements. Nature 460, 627–631, doi:nature08209 [pii] 10.1038/nature08209 (2009).

6 Fung, C. W. et al. Cell fate determining molecular switches and signaling pathways in Pax7-expressing somitic mesoderm. Cell Discov 8, 61, doi:10.1038/s41421-022-00407-0 (2022).

7 Grimaldi, A., Comai, G., Mella, S. & Tajbakhsh, S. Identification of bipotent progenitors that give rise to myogenic and connective tissues in mouse. Elife 11, doi:10.7554/eLife.70235 (2022).

8 Esteves de Lima, J. et al. Unexpected contribution of fibroblasts to muscle lineage as a mechanism for limb muscle patterning. Nature communications 12, 3851, doi:10.1038/s41467-021-24157-x (2021).

9 Yaseen, W. et al. Fibroblast fusion to the muscle fiber regulates myotendinous junction formation. Nature communications 12, 3852, doi:10.1038/s41467-021-24159-9 (2021).

10 Jagot, S., Babarit, C., Sabin, N. & Gabillard, P. C. Distinct muscle stem cell fates governing hyperplasia and hypertrophy muscle growth in fish. bioRxiv, doi: 10.64898/2026.01.28.702282 (2026).

11 Sambasivan, R. et al. Pax7-expressing satellite cells are indispensable for adult skeletal muscle regeneration. Development 138, 3647–3656, doi:10.1242/dev.067587 (2011).

12 Lepper, C., Partridge, T. A. & Fan, C. M. An absolute requirement for Pax7-positive satellite cells in acute injury-induced skeletal muscle regeneration. Development 138, 3639–3646, doi:10.1242/dev.067595 (2011).

13 McCarthy, J. J. et al. Effective fiber hypertrophy in satellite cell-depleted skeletal muscle. Development 138, 3657–3666, doi:10.1242/dev.068858 (2011).

14 Murphy, M. M., Lawson, J. A., Mathew, S. J., Hutcheson, D. A. & Kardon, G. Satellite cells, connective tissue fibroblasts and their interactions are crucial for muscle regeneration. Development 138, 3625–3637, doi:10.1242/dev.064162 (2011).

15 Biressi, S., Miyabara, E. H., Gopinath, S. D., Carlig, P. M. & Rando, T. A. A Wnt-TGFbeta2 axis induces a fibrogenic program in muscle stem cells from dystrophic mice. Sci Transl Med 6, 267ra176, doi:10.1126/scitranslmed.3008411 (2014).

16 Pessina, P. et al. Fibrogenic Cell Plasticity Blunts Tissue Regeneration and Aggravates Muscular Dystrophy. Stem Cell Reports 4, 1046–1060, doi:10.1016/j.stemcr.2015.04.007 (2015).

17 Roberts, E. W. et al. Depletion of stromal cells expressing fibroblast activation proteinalpha from skeletal muscle and bone marrow results in cachexia and anemia. The Journal of experimental medicine 210, 1137–1151, doi:10.1084/jem.20122344 (2013).

18 Wosczyna, M. N. et al. Mesenchymal Stromal Cells Are Required for Regeneration and Homeostatic Maintenance of Skeletal Muscle. Cell Rep 27, 2029–2035 e2025, doi:10.1016/j.celrep.2019.04.074 (2019).

19 Lilja, A. M. et al. Clonal analysis of Notch1-expressing cells reveals the existence of unipotent stem cells that retain long-term plasticity in the embryonic mammary gland. Nat Cell Biol 20, 677–687, doi:10.1038/s41556-018-0108-1 (2018).

20 Lepper, C. & Fan, C. M. Inducible lineage tracing of Pax7-descendant cells reveals embryonic origin of adult satellite cells. Genesis 48, 424–436, doi:10.1002/dvg.20630 (2010).

21 Machado, L. et al. In Situ Fixation Redefines Quiescence and Early Activation of Skeletal Muscle Stem Cells. Cell Rep 21, 1982–1993, doi:10.1016/j.celrep.2017.10.080 (2017).

22 Machado, L. et al. Tissue damage induces a conserved stress response that initiates quiescent muscle stem cell activation. Cell stem cell 21, 1934–5909, doi:10.1016/j.stem.2021.01.017 (2021).

23 Machado, L., Relaix, F. & Mourikis, P. Stress relief: emerging methods to mitigate dissociation-induced artefacts. Trends Cell Biol 31, 888–897, doi:10.1016/j.tcb.2021.05.004 (2021).

24 Dos Santos, M. D. et al. Extraction and sequencing of single nuclei from murine skeletal muscles. STAR Protoc 2, 100694, doi:10.1016/j.xpro.2021.100694 (2021).

25 White, R. B., Bierinx, A. S., Gnocchi, V. F. & Zammit, P. S. Dynamics of muscle fibre growth during postnatal mouse development. BMC developmental biology 10, 21, doi:10.1186/1471-213X-10-21 (2010).

26 Bachman, J. F. et al. Prepubertal skeletal muscle growth requires Pax7-expressing satellite cell-derived myonuclear contribution. Development 145, doi:10.1242/dev.167197 (2018).

27 Coren, L., Zaffryar-Eilot, S., Odeh, A., Kaganovsky, A. & Hasson, P. Fibroblast diversification is an embryonic process dependent on muscle contraction. Cell Rep 43, 115034, doi:10.1016/j.celrep.2024.115034 (2024).

28 Buechler, M. B. et al. Cross-tissue organization of the fibroblast lineage. Nature 593, 575–579, doi:10.1038/s41586-021-03549-5 (2021).

29 Artavanis-Tsakonas, S., Matsuno, K. & Fortini, M. E. Notch signaling. Science 268, 225–232 (1995).

30 Mourikis, P. et al. A critical requirement for notch signaling in maintenance of the quiescent skeletal muscle stem cell state. Stem Cells 30, 243–252, doi:10.1002/stem.775 (2012).

31 Gioftsidi, S., Relaix, F. & Mourikis, P. The Notch signaling network in muscle stem cells during development, homeostasis, and disease. Skeletal muscle 12, 9, doi:10.1186/s13395-022-00293-w (2022).

32 Mourikis, P., Gopalakrishnan, S., Sambasivan, R. & Tajbakhsh, S. Cell-autonomous Notch activity maintains the temporal specification potential of skeletal muscle stem cells. Development 139, 4536–4548, doi:10.1242/dev.084756 (2012).

33 Latroche, C., Weiss-Gayet, M., Gitiaux, C. & Chazaud, B. Cell sorting of various cell types from mouse and human skeletal muscle. Methods 134-135, 50–55, doi:10.1016/j.ymeth.2017.12.013 (2018).

34 Hutcheson, D. A., Zhao, J., Merrell, A., Haldar, M. & Kardon, G. Embryonic and fetal limb myogenic cells are derived from developmentally distinct progenitors and have different requirements for beta-catenin. Genes & development 23, 997–1013, doi:10.1101/gad.1769009 (2009).

35 Murphy, M. & Kardon, G. Origin of vertebrate limb muscle: the role of progenitor and myoblast populations. Curr Top Dev Biol 96, 1–32, doi:10.1016/B978-0-12-385940-2.00001-2 (2011).

36 Sefton, E. M. & Kardon, G. Connecting muscle development, birth defects, and evolution: An essential role for muscle connective tissue. Curr Top Dev Biol 132, 137–176, doi:10.1016/bs.ctdb.2018.12.004 (2019).

37 Chevallier, A., Kieny, M. & Mauger, A. Limb-somite relationship: origin of the limb musculature. J Embryol Exp Morphol 41, 245–258 (1977).

38 Christ, B., Jacob, H. J. & Jacob, M. Experimental analysis of the origin of the wing musculature in avian embryos. Anat Embryol (Berl) 150, 171–186, doi:10.1007/BF00316649 (1977).

39 Kardon, G., Harfe, B. D. & Tabin, C. J. A Tcf4-positive mesodermal population provides a prepattern for vertebrate limb muscle patterning. Developmental cell 5, 937–944, doi:10.1016/s1534-5807(03)00360-5 (2003).

40 Mathew, S. J. et al. Connective tissue fibroblasts and Tcf4 regulate myogenesis. Development 138, 371–384, doi:10.1242/dev.057463 (2011).

41 Vallecillo-Garcia, P. et al. Odd skipped-related 1 identifies a population of embryonic fibro-adipogenic progenitors regulating myogenesis during limb development. Nature communications 8, 1218, doi:10.1038/s41467-017-01120-3 (2017).

42 Uezumi, A., Fukada, S., Yamamoto, N., Takeda, S. & Tsuchida, K. Mesenchymal progenitors distinct from satellite cells contribute to ectopic fat cell formation in skeletal muscle. Nat Cell Biol 12, 143–152, doi:ncb2014 [pii] 10.1038/ncb2014 (2010).

43 Kardon, G., Campbell, J. K. & Tabin, C. J. Local extrinsic signals determine muscle and endothelial cell fate and patterning in the vertebrate limb. Developmental cell 3, 533–545, doi:10.1016/s1534-5807(02)00291-5 (2002).

44 Ben-Yair, R. & Kalcheim, C. Notch and bone morphogenetic protein differentially act on dermomyotome cells to generate endothelium, smooth, and striated muscle. J Cell Biol 180, 607–618, doi:10.1083/jcb.200707206 (2008).

45 Mayeuf-Louchart, A. et al. Notch regulation of myogenic versus endothelial fates of cells that migrate from the somite to the limb. Proceedings of the National Academy of Sciences of the United States of America 111, 8844–8849, doi:10.1073/pnas.1407606111 (2014).

46 Luth, E. S. et al. Bone marrow side population cells are enriched for progenitors capable of myogenic differentiation. Journal of cell science 121, 1426–1434, doi:10.1242/jcs.021675 (2008).

47 Marinkovic, M. et al. Fibro-adipogenic progenitors of dystrophic mice are insensitive to NOTCH regulation of adipogenesis. Life Sci Alliance 2, doi:10.26508/lsa.201900437 (2019).

48 Giordani, L. et al. High-Dimensional Single-Cell Cartography Reveals Novel Skeletal Muscle-Resident Cell Populations. Mol Cell 74, 609–621 e606, doi:10.1016/j.molcel.2019.02.026 (2019).

49 Merrick, D. et al. Identification of a mesenchymal progenitor cell hierarchy in adipose tissue. Science 364, doi:10.1126/science.aav2501 (2019).

50 Scott, R. W., Arostegui, M., Schweitzer, R., Rossi, F. M. V. & Underhill, T. M. Hic1 Defines Quiescent Mesenchymal Progenitor Subpopulations with Distinct Functions and Fates in Skeletal Muscle Regeneration. Cell stem cell 25, 797–813 e799, doi:10.1016/j.stem.2019.11.004 (2019).

51 Oprescu, S. N., Yue, F., Qiu, J., Brito, L. F. & Kuang, S. Temporal Dynamics and Heterogeneity of Cell Populations during Skeletal Muscle Regeneration. iScience 23, 100993, doi:10.1016/j.isci.2020.100993 (2020).

52 Leinroth, A. P. et al. Identification of distinct non-myogenic skeletal-muscle-resident mesenchymal cell populations. Cell Rep 39, 110785, doi:10.1016/j.celrep.2022.110785 (2022).

53 Giuliani, G. et al. SCA-1 micro-heterogeneity in the fate decision of dystrophic fibro/adipogenic progenitors. Cell Death Dis 12, 122, doi:10.1038/s41419-021-03408-1 (2021).

54 Yao, L. et al. Gli1 Defines a Subset of Fibro-adipogenic Progenitors that Promote Skeletal Muscle Regeneration With Less Fat Accumulation. J Bone Miner Res 36, 1159–1173, doi:10.1002/jbmr.4265 (2021).

55 Brack, A. S. et al. Increased Wnt signaling during aging alters muscle stem cell fate and increases fibrosis. Science 317, 807–810, doi:10.1126/science.1144090 (2007).

56 De Angelis, L. et al. Skeletal myogenic progenitors originating from embryonic dorsal aorta coexpress endothelial and myogenic markers and contribute to postnatal muscle growth and regeneration. J Cell Biol 147, 869–878, doi:10.1083/jcb.147.4.869 (1999).

57 Minasi, M. G. et al. The meso-angioblast: a multipotent, self-renewing cell that originates from the dorsal aorta and differentiates into most mesodermal tissues. Development 129, 2773–2783, doi:10.1242/dev.129.11.2773 (2002).

58 Le Grand, F. et al. Endothelial cells within embryonic skeletal muscles: a potential source of myogenic progenitors. Exp Cell Res 301, 232–241, doi:10.1016/j.yexcr.2004.07.028 (2004).

59 Liu, N. et al. A Twist2-dependent progenitor cell contributes to adult skeletal muscle. Nat Cell Biol 19, 202–213, doi:10.1038/ncb3477 (2017).

60 Flynn, C. G. K. et al. Hox11-expressing interstitial cells contribute to adult skeletal muscle at homeostasis. Development 150, doi:10.1242/dev.201026 (2023).

61 Sambasivan, R. et al. Distinct regulatory cascades govern extraocular and pharyngeal arch muscle progenitor cell fates. Developmental cell 16, 810–821, doi:10.1016/j.devcel.2009.05.008 (2009).

62 Murtaugh, L. C., Stanger, B. Z., Kwan, K. M. & Melton, D. A. Notch signaling controls multiple steps of pancreatic differentiation. Proceedings of the National Academy of Sciences of the United States of America 100, 14920–14925, doi:10.1073/pnas.2436557100 (2003).

63 Hamilton, T. G., Klinghoffer, R. A., Corrin, P. D. & Soriano, P. Evolutionary divergence of platelet-derived growth factor alpha receptor signaling mechanisms. Molecular and cellular biology 23, 4013–4025, doi:10.1128/MCB.23.11.4013-4025.2003 (2003).

64 Li, Y. et al. Molecular layer perforant path-associated cells contribute to feed-forward inhibition in the adult dentate gyrus. Proceedings of the National Academy of Sciences of the United States of America 110, 9106–9111, doi:10.1073/pnas.1306912110 (2013).

65 Muzumdar, M. D., Tasic, B., Miyamichi, K., Li, L. & Luo, L. A global double-fluorescent Cre reporter mouse. Genesis 45, 593–605, doi:10.1002/dvg.20335 (2007).

66 Madisen, L. et al. A robust and high-throughput Cre reporting and characterization system for the whole mouse brain. Nat Neurosci 13, 133–140, doi:10.1038/nn.2467 (2010).

67 Lizen, B., Claus, M., Jeannotte, L., Rijli, F. M. & Gofflot, F. Perinatal induction of Cre recombination with tamoxifen. Transgenic Res 24, 1065–1077, doi:10.1007/s11248-015-9905-5 (2015).

