## Supplemental Figures for "Postnatal Pax7-expressing limb cells are multipotent and generate non-myogenic lineages that persist into adulthood"

### Figure S1

A

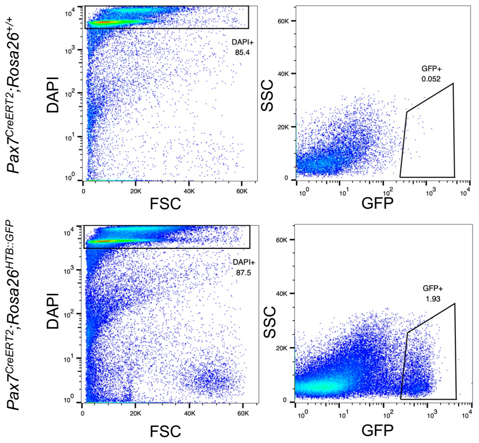


B

| single-nucleusRNA-seq | | | | | |
|---|---|---|---|---|---|
| Age | Sequenced nuclei | Nuclei after filtering | Mean reads per nucleus | Mean genes per nucleus | Type of experiment |
| P3 | 6 121 | 4 275 | 43K | 1 822 | Pax7 lineage |
| P3 | 8 130 | 7 398 | 29K | 1 646 | Whole muscle |
| P10 | 3 600 | 3 205 | 164K | 1 471 | Pax7 lineage |
| P22 | 3 923 | 2 856 | 133K | 1 092 | Pax7 lineage |
| Adult | 10 081 | 8 878 | 80K | 1 145 | Pax7 lineage |

C

| Cluster | WP3 | P3 | P10 | P22 |
|---|---|---|---|---|
| **0.FAPs (Smoc2+)** | 57.7 | 12.6 | 5.8 | 2.7 |
| **1.qMuSCs** | 2.6 | 33.6 | 30.5 | 28.9 |
| **2.MyoNuc** | 8.3 | 11 | 26.7 | 44.7 |
| **3.pMuSCs** | 0.9 | 16.2 | 12.3 | 6.1 |
| **4.Myocytes** | 0.6 | 8.5 | 15.5 | 10.6 |
| **5.Tenocytes** | 9.4 | 2.9 | 1.5 | 1.1 |
| **6.Endothelial cells** | 4.3 | 3.4 | 1.9 | 1.5 |
| **7.Macrophages** | 4.4 | 2.9 | 1.5 | 0.8 |
| **8.MyoFAPs** | 2.7 | 2.6 | 1.1 | 0.4 |
| **9.Lymphocytes** | 1.5 | 1.8 | 1 | 2.2 |
| **10.FAPs (Tenm2+)** | 2.2 | 0.9 | 0.5 | 0.1 |
| **11.Mural cells** | 2.1 | 0.8 | 0.8 | 0.3 |
| **12.Mesenchymal cells** | 1.4 | 1.7 | 0.3 | 0.1 |
| **13.Schwann cells** | 1.2 | 0.4 | 0.2 | 0.3 |
| **14.Adipocytes** | 0.7 | 0.6 | 0.5 | 0.2 |

D

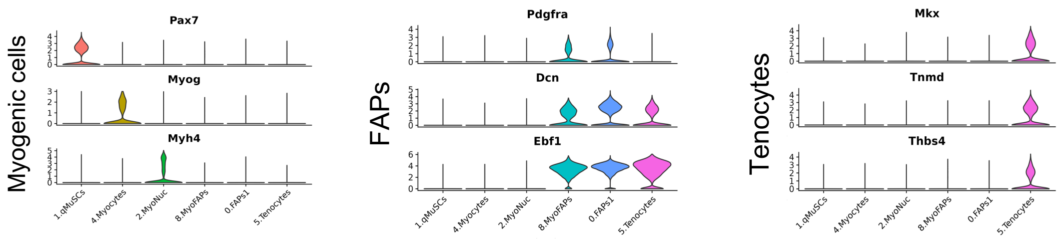


E

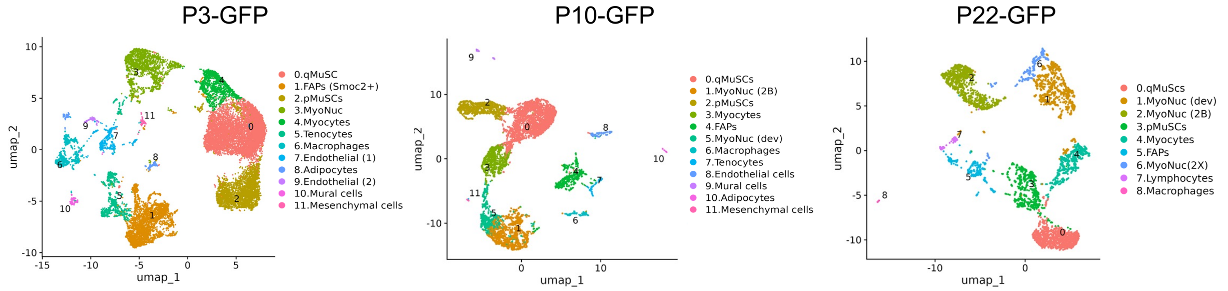


F

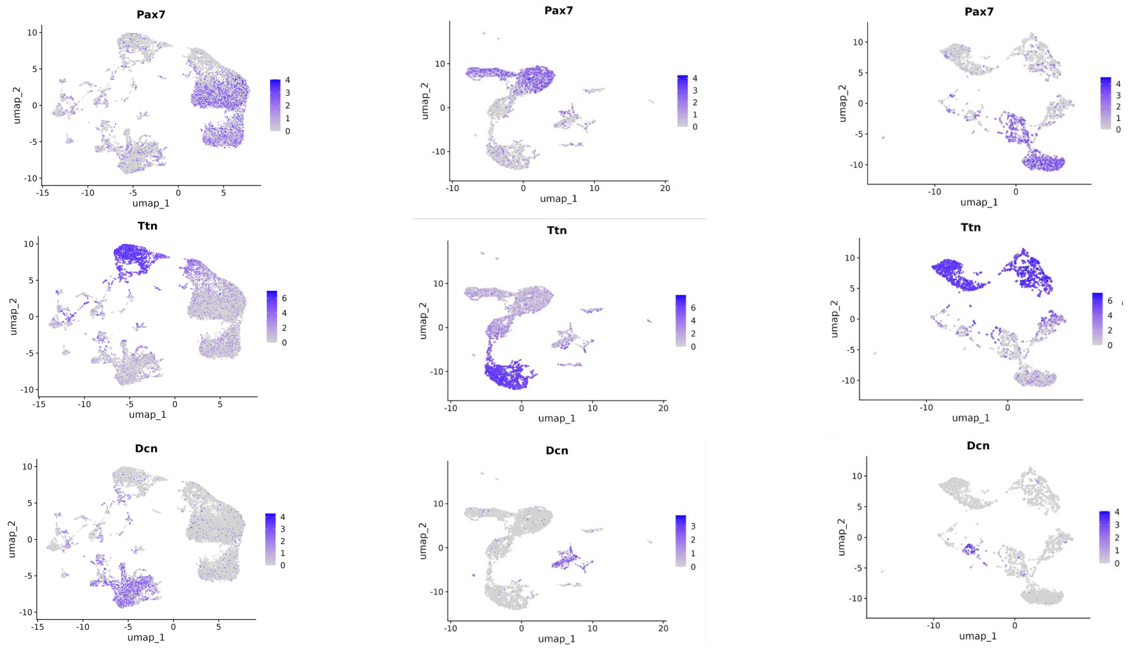

**Figure S1. Single-nucleus transcriptomic profiling and cluster annotation of Pax7⁺ lineage cells in postnatal muscle.**

**(A)** FACS gating for the *Pax7*$^{CreERT2}$*;Rosa26*$^{HTB::GFP}$ mice. Nuclei were selected based on 4,6-diamidino-2-phenylindole (DAPI), followed by selection for GFP for the H2B::GFP marked nuclei. All mice, including controls, were injected twice with tamoxifen, as shown in figure 1A.

**(B)** Table with quantitative information on the different snRNA-seq experiments.

**(C)** Table of absolute cell numbers per cluster in each sample, corresponding to the integrated UMAP in figure 1B (including Pax7-lineage cells from hindlimb muscles collected at P3, P10, and P22 (48 h chase), as well as whole hindlimb muscles from P3 neonatal mice).

**(D)** Violin plots of gene expression for markers in clusters from the integrated UMAP, annotated as myogenic cells (*Pax7*, *Myog*, *Myh4*), main FAPs cluster 0 (*Pdgfra*, *Dcn*, *Ebf1*), and tenocytes (*Mkx*, *Tnmd*, *Thbs4*).

**(E)** UMAP with individual analysis of data at each time point.

**(F)** Feature plots showing the expression levels of *Pax7*, *Ttn*, and *Dcn* at P3-GFP, P10-GFP, and P22-GFP.

### Figure S2

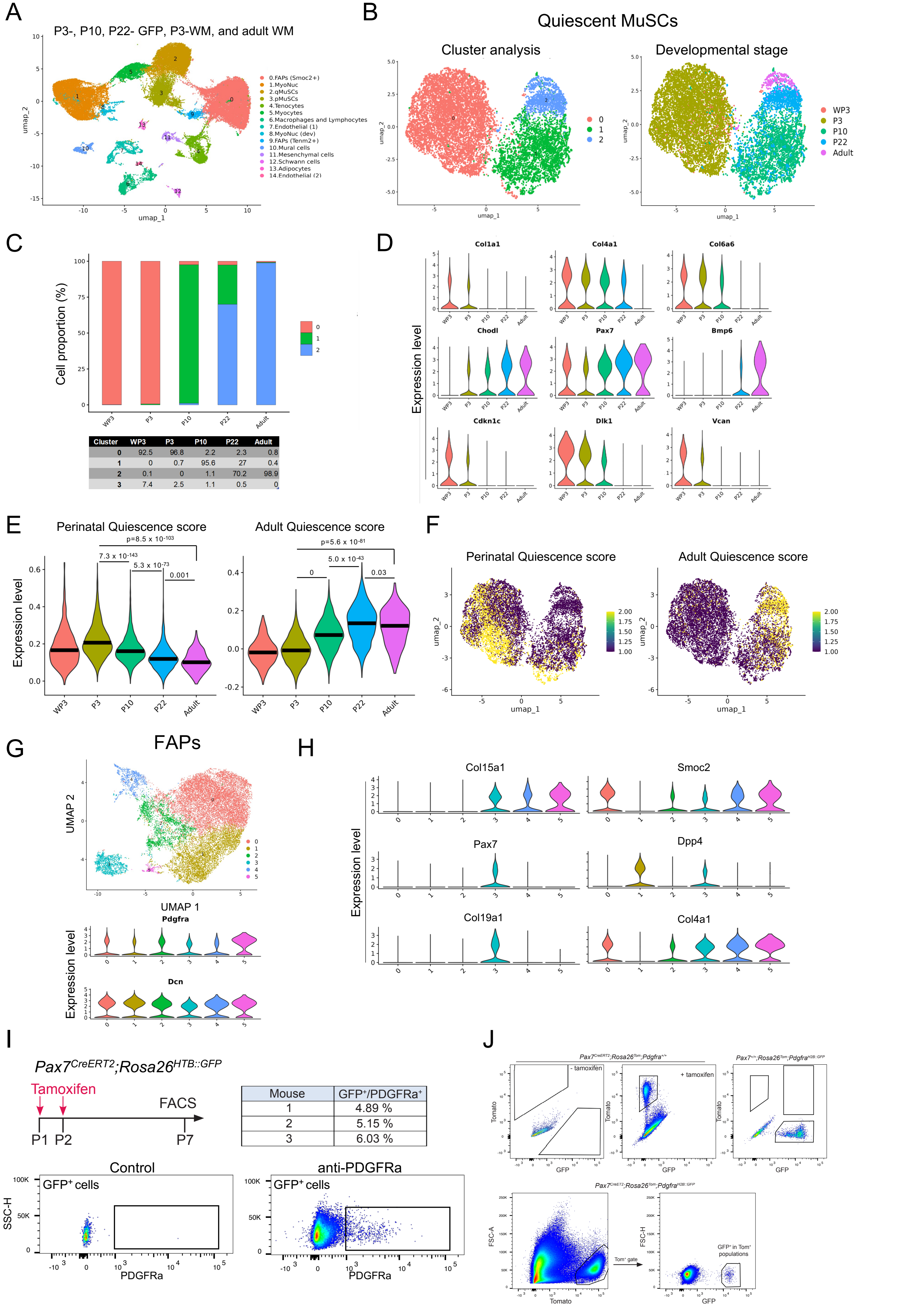


| Cluster | WP3 | P3 | P10 | P22 | Adult |
|---|---|---|---|---|---|
| 0 | 92.5 | 96.8 | 2.2 | 2.3 | 0.8 |
| 1 | 0 | 0.7 | 95.6 | 27 | 0.4 |
| 2 | 0.1 | 0 | 1.1 | 70.2 | 98.9 |
| 3 | 7.4 | 2.5 | 1.1 | 0.5 | 0 |

| Mouse | $GFP^+/PDGFRa^+$ |
|---|---|
| 1 | 4.89 % |
| 2 | 5.15 % |
| 3 | 6.03 % |

**Figure S2 Comparison of quiescent MuSCs and Pax7-derived FAPs across postnatal and adult muscles**

**(A)** Integrated UMAP of snRNA-seq experiments of Pax7 lineage at P3, P10, and P22, whole hindlimb muscles from P3, and TA muscle from 8-week-old mice (adult dataset from Machado et al., 2021 (22).

**(B)** *Left*: Higher definition reclustering of the quiescent MuSCs population extracted from the integrated UMAP (Figure 1B). *Right*: Distribution of the quiescent MuSCs from P3, P10 and P22 Pax7-lineage traced datasets, whole hindlimb muscles from P3 and 8-week-old adult whole TA muscle.

**(C)** Stacked bar graph representing the proportion of cells in each cluster of quiescent MuSCs at different developmental stages. Table corresponding to percentages of total cells per cluster.

**(D)** Violin plots depicting cluster-wise expression of MuSCs quiescence markers (*Chodl, Pax7, Bmp6*), extracellular matrix-associated genes (*Col1a1, Col4a1, Col6a6, Vcan*), *Dlk1, and* cell cycle gene *Cdkn1c (p57).*

(**E**) Module scores for Perinatal and Adult Quiescence in quiescent MuSCs (qMuSCs) across developmental stages. The Perinatal Quiescence module was defined as the median expression of genes significantly over-expressed in P3-GFP q-MuSCs compared to proliferating MuSCs (prolifMuSCs; see UMAP in Fig. S1E), applying a threshold of >0.25 log fold induction and padj < 0.001. The Adult Quiescence module was calculated as the median expression of genes significantly over-expressed in T0 qMuSCs versus T3 activated-MuSCs (Machado et al., 2017), using a threshold of >0.25 log fold induction and an expression level >5,000 RPKM. Line indicates the median value. Statistical significance was determined by the Kruskal-Wallis test (Perinatal; $H$ = 2043, p < 0.001, Adult; $H$ = 3414, p < 0.001). Pairwise comparisons were performed using Dunn's test with Benjamini-Hochberg correction.

(**F**) Projection of Perinatal and Adult Quiescence module scores onto UMAP coordinates, visualized by color intensity.

**(G)** UMAP of the FAPs clusters subset from Figure 1B. Violin plots showing the expression of *Pdgfra* and *Dcn* in each cluster.

**(H)** Violin plots showing the expression of myogenic markers (*Pax7*, *Col19a1*) and fibrogenic markers (*Col15a1*, *Smoc2*, *Dpp4*, *Col4a1*) across clusters.

**Figure S2 –continued**

**(I)** *Pax7*$^{CreERT2}$*;Rosa26*$^{HTB::GFP}$ mice induced at P1 and P2 and hindlimb muscle collected at P7. Cells were stained for PDGFRa and analyzed by FACS. Non stained muscle bulk preparation was used as control. The table shows the percentage of PDGFRa+ cells in the GFP+ population (n=3).

**(J)** Gating strategy for the identification and quantification of double positive, Tom+/GFP+ cells from *Pax7*$^{CreERT2/+}$*; Rosa26* $^{Tom/+}$*;Pdgfra*$^{H2B::GFP/+}$ muscle preparations, induced at P1 and P2 and killed at P3. *Upper panels*: Hindlimb muscles from non-induced or tamoxifen-induced mice to test for aberrant Tomato induction and leakiness (*Pax7*$^{CreERT2/+}$*; Rosa26* $^{Tom/+}$ mice). *Right*: Hindlimb muscles from *Pdgfra*$^{H2B::GFP/+}$ mice at P3. *Lower panels*: *Pax7*$^{CreERT2/+}$*; Rosa26* $^{Tom/+}$*;Pdgfra*$^{H2B::GFP/+}$ muscle preparations, induced with tamoxifen at P1 and P2 and collected at P3.

### Figure S3

A

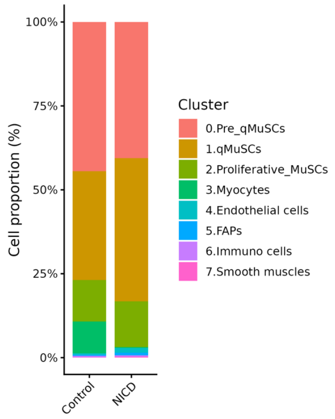


| Cluster | Control | NICD |
|---|---|---|
| **0.Pre_qMuSCs** | 44.5 | 40.5 |
| **1.qMuSCs** | 32.4 | 42.7 |
| **2.pMuSCs** | 12.4 | 13.6 |
| **3.Myocytes** | 9.4 | 0.2 |
| **4.Endothelial cells** | 0.4 | 1.4 |
| **5.FAPs** | 0.4 | 0.7 |
| **6.Immuno cells** | 0.4 | 0.5 |
| **7.Smooth muscles** | 0.1 | 0.2 |

B

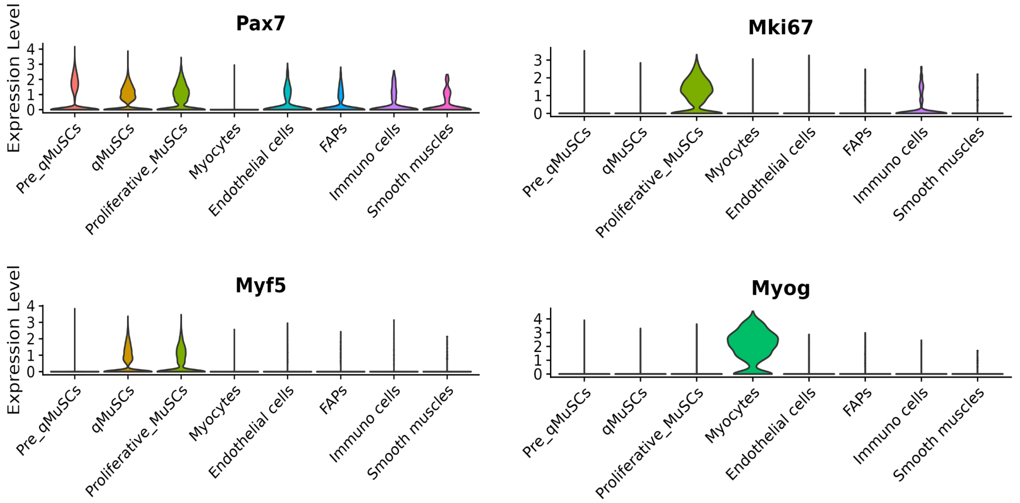

**Figure S3. Impact of Notch activation on Pax7 lineage cells**

**(A)** Left: Stacked bar graph representing the proportion of cells in each cluster of the integrated UMAP (Fig. 3A) in control and Pax7-NICD datasets. Right: Proportion of total cells in each cluster (%).

**(B)** Violin plots of myogenic markers (*Pax7*, *Myf5*, *Myog*) and proliferation marker *Mki67* across clusters.

### Figure S4

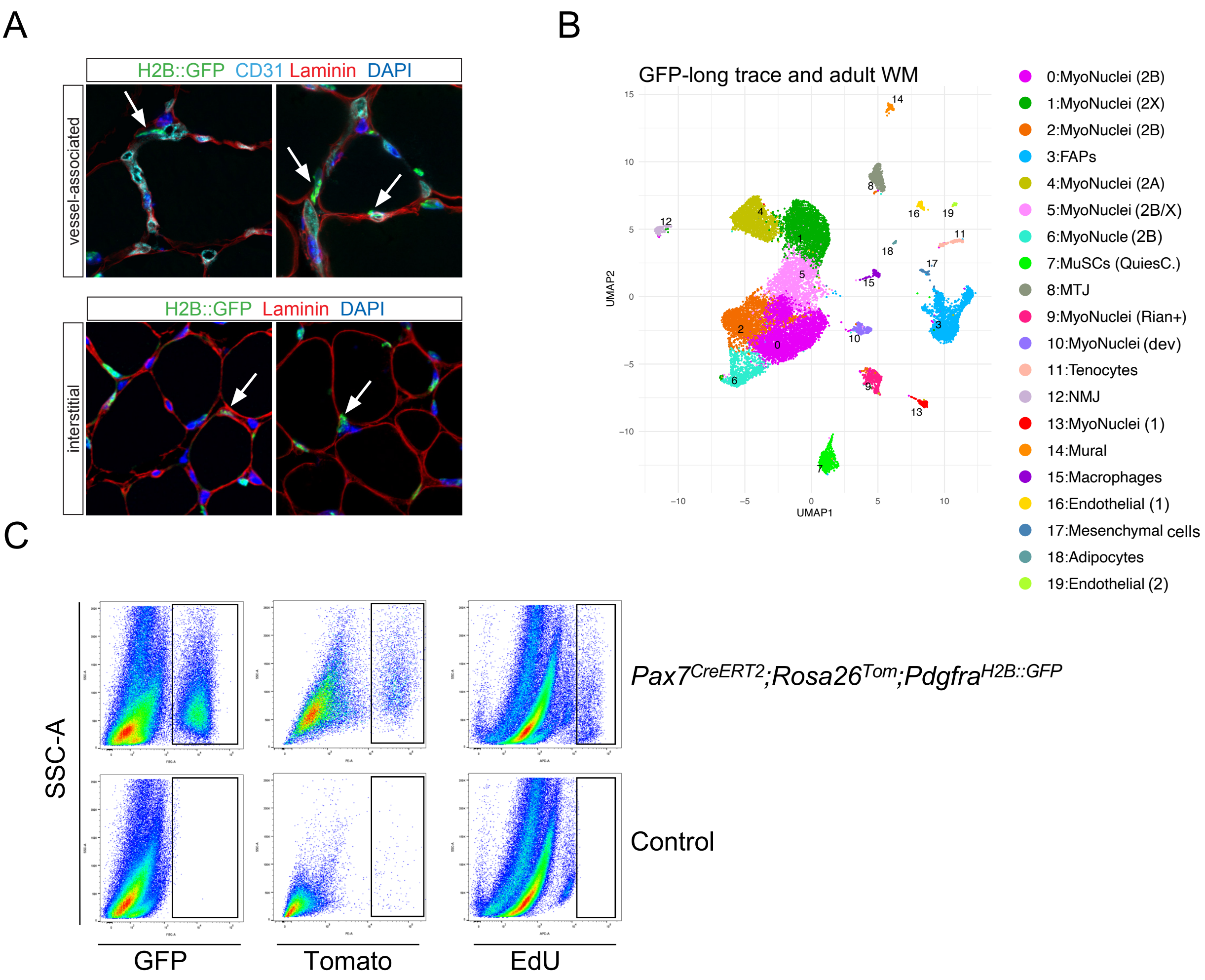


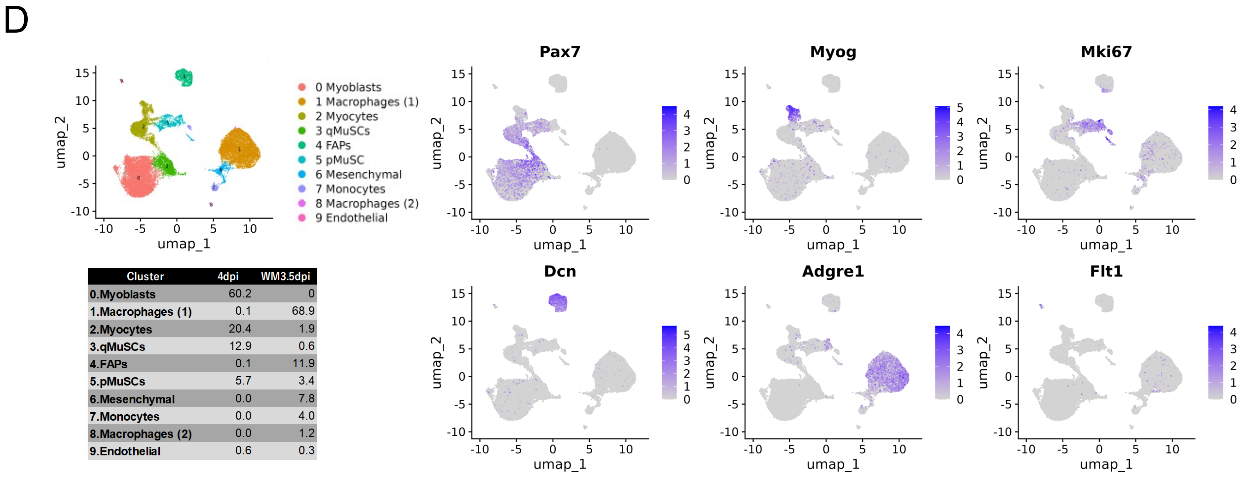


| Cluster | 4dpi | WM3.5dpi |
|---|---|---|
| 0.Myoblasts | 60.2 | 0 |
| 1.Macrophages (1) | 0.1 | 68.9 |
| 2.Myocytes | 20.4 | 1.9 |
| 3.qMuSCs | 12.9 | 0.6 |
| 4.FAPs | 0.1 | 11.9 |
| 5.pMuSCs | 5.7 | 3.4 |
| 6.Mesenchymal | 0.0 | 7.8 |
| 7.Monocytes | 0.0 | 4.0 |
| 8.Macrophages (2) | 0.0 | 1.2 |
| 9.Endothelial | 0.6 | 0.3 |

**Figure S4. Localization, transcriptomic profiling, and injury-induced proliferation of adult Pax7 lineage cells.**

**(A)** Interstitial and vessel-associated GFP+ cells in adult *Pax7*$^{CreERT2}$;*Rosa26*$^{HTB::GFP}$ mice induced with tamoxifen at P1-P4. TA sections were stained for GFP/Laminin for interstitial cells (2-month old mice) or for GFP/Laminin/CD31 (5.5-week-old mice) to identify vessel-associated cells. Arrows point to Pax7 lineage-traced GFP+ cells.

**(B)** Detailed annotation of snRNA-seq data of long lineage tracing also shown in Figure. 4E. *Pax7*$^{CreERT2}$;*Rosa26*$^{HTB::GFP}$ mice were induced at P1-P4 and sacrificed after 2 months. (DAPI+GFP+ nuclei from hindlimb muscles, n=4 mice pooled).

**(C)** Gating strategy for the identification and quantification of Tom$^{+}$/GFP$^{+}$/EdU$^{+}$ cells from muscle preparations of *Pax7*$^{CreERT2}$;*Rosa26*$^{Tom/+}$;*Pdgfra*$^{H2B::GFP/+}$ mice. Quantification of each cell population was performed based on GFP-positive cells (left panel), Tomato-positive cells (middle panel), and EdU-positive cells (Far-red) (right panel), with gating thresholds determined by comparison to control mouse samples.

**(D)** Upper left: Integrated UMAP visualization of scRNA-seq data, including FACS-isolated Pax7$^{+}$ lineage traced cells (*Pax7*$^{CreERT2}$;*Rosa26*$^{HTB::GFP}$) collected at 4 days post injury (dpi), integrated with the 3.5 dpi whole TA muscle dataset from Oprescu et al., 2020 (51). Lower left: Table showing percentage of cell numbers per cluster in each sample, corresponding to the integrated UMAP. Right panels: Feature plots displaying expression profiles of myogenic and niche-related marker genes, including *Pax7*, *Myog*, *Mki67*, *Dcn* (FAPs), *Adgre1* (macrophages), and *Flt1* (endothelial cells).
